# High affinity modified ACE2 receptors protect from SARS-CoV-2 infection in hamsters

**DOI:** 10.1101/2020.09.16.299891

**Authors:** Yusuke Higuchi, Tatsuya Suzuki, Takao Arimori, Nariko Ikemura, Emiko Mihara, Yuhei Kirita, Eriko Ohgitani, Osam Mazda, Daisuke Motooka, Shota Nakamura, Yusuke Sakai, Yumi Itoh, Fuminori Sugihara, Yoshiharu Matsuura, Satoaki Matoba, Toru Okamoto, Junichi Takagi, Atsushi Hoshino

## Abstract

The SARS-CoV-2 spike protein binds to the human angiotensin-converting enzyme 2 (ACE2) receptor via receptor binding domain (RBD) to enter into the cell and inhibiting this interaction is a main approach to inhibit SARS-CoV-2 infection. We engineered ACE2 to enhance the affinity with directed evolution in 293T cells. Three cycles of random mutation and cell sorting achieved 100-fold higher affinity to RBD than wild-type ACE2. The extracellular domain of modified ACE2 fused to the human IgG1-Fc region had stable structure and neutralized SARS-CoV-2 without the emergence of mutational escape. Therapeutic administration protected hamsters from SARS-CoV-2 infection, decreasing lung virus titers and pathology. Engineering ACE2 decoy receptors with human cell-based directed evolution is a promising approach to develop a SARS-CoV-2 neutralizing drug that has affinity comparable to monoclonal antibodies yet displaying resistance to escape mutations of virus.

**One Sentence Summary:** Engineered ACE2 decoy receptor has a therapeutic potential against COVID-19 without viral escape mutation.

## Main text

Coronavirus disease 2019 (COVID-19) has spread across the world as a tremendous pandemic and presented an unprecedented challenge to human society. The causative agent of COVID-19, SARS-CoV-2 is a single-stranded positive-strand RNA virus that belongs to lineage B, clade 1 of the betacoronavirus genus(*1-3*). The virus binds to host cells through its trimeric spike glycoprotein composed of two subunits; S1 is responsible for receptor binding and S2 for membrane fusion(*4*). Angiotensin-converting enzyme 2 (ACE2) is lineage B clade 1 specific receptor including SARS-CoV-2 (*3*). The receptor binding domain (RBD) of S1 subunit directly binds ACE2, therefore, it is the most important targeting site to inhibit viral infection. In fact, the RBD is the common binding site of effective neutralizing antibodies identified from convalescent patients (*5-7*).

RNA viruses such as SARS-CoV-2 have high mutation rates (*8*), which are correlated with high evolvability including the acquisition of anti-viral drug resistance. Accumulating evidence demonstrated that monoclonal antibodies isolated from convalescent COVID-19 patients have high potency in neutralizing viruses. However, mutations in the spike gene can lead to the SARS-CoV-2 adaptation to such neutralizing antibodies. In the replicating SARS-CoV-2 pseudovirus culture experiment, escape mutation was observed against monoclonal antibody as early as in the first passage (*9*) and virus evasion was seen even against the polyclonal convalescent plasma (*10*). Notably, some mutations identified in *in vitro* replicating culture experiment are present in natural population according to the database (*10*), indicating that selection and expansion of escape mutants can occur naturally not only in the therapeutic use of monoclonal antibodies, but in the vaccination that was recently approved to use in the mRNA vaccine platform.

Similar to anti-RBD antibodies, extracellular domain of ACE2, soluble ACE2 (sACE2), can also be used to neutralize SARS-CoV-2 as a decoy receptor. The therapeutic potency was confirmed using human organoid (*11*), and now Apeiron Biologics conducts European phase II clinical trial of recombinant sACE2 against COVID-19 (*12*). In addition, fusing sACE2 to the Fc region of the human IgG1 has been shown to enhance neutralization capacity (*13*) as well as to improve the pharmacokinetics to the level of IgG in mice (*14*). Most importantly, sACE2 has a great advantage over antibodies due to the resistance to the escape mutation. The virus with escape mutation from sACE2 should have limited binding affinity to cell surface native ACE2 receptors, leading to a diminished or eliminated virulence. Unfortunately, many reports, including our current study, have revealed that the binding affinity of wild-type (WT) sACE2 to the SARS-CoV-2 spike RBD is much weaker (K_D_ ∼20 nM) than that of clinical grade antibodies (*4, 13, 15, 16*). Thus, the therapeutic potential of the WT sACE2 as a neutralizing agent against SARS-CoV-2 is uncertain.

Here we conducted human cell-based directed evolution to improve the binding affinity of ACE2 to the spike RBD with the combination of surface display of mutagenized library and fluorescence-activated cell sorting (FACS). Among various host cells, the budding yeast is popular in directed evolution due to the efficient surface display and greater molecular diversity (*17*). However, posttranslational modifications such as glycosylation are substantially different between yeast and mammalian cells, which could alter the protein activity including binding affinity (*18*). In consideration of future drug development in mammalian cells, we developed the screening system based on 293T cells. The protease domain (PD) of ACE2 is known to harbor the interface to viral spike protein, located in the top-middle part of ACE2 ectodomain. In this study, ACE2 residues 18-102 and 272-409, referred to as PD1 and PD2, respectively, were mutagenized independently. Synthetic signal sequence and HA tag were appended and restriction sites were introduced in both sides of PD1 and PD2 by optimizing codon (Fig. 1A). We used error-prone PCR to mutagenize the protease domain of ACE2 with an average of about one amino acid mutation per 100 bp, then inserted the fragment into the introduced restriction site. The reaction sample was transformed to competent cell, generating a library of ∼10^5^ mutants. Mutant plasmid library was packaged into lentivirus, followed by expression in human 293T cells in less than 0.3 MOI (multiplicity of infection) to yield no more than one mutant ACE2 per cell. Cells were incubated with recombinant RBD of SARS-CoV-2 spike protein fused to superfolder GFP (sfGFP; Fig. 1B). We confirmed the level of bound RBD-sfGFP and surface expression levels of HA-tagged ACE2 with Alexa Fluor 647 in two-dimensional display of flow cytometry. Top 0.05 % cells showing higher binding relative to expression level were harvested from ∼5 × 10^7^ cells by FACS. To exclude the structurally unstable mutants, cells with preserved signal of surface ACE2 were gated. Genomic DNA was extracted from collected cells and mutagenized again to proceed to the next cycle of screening (Fig. 1C).

**Fig 1.**
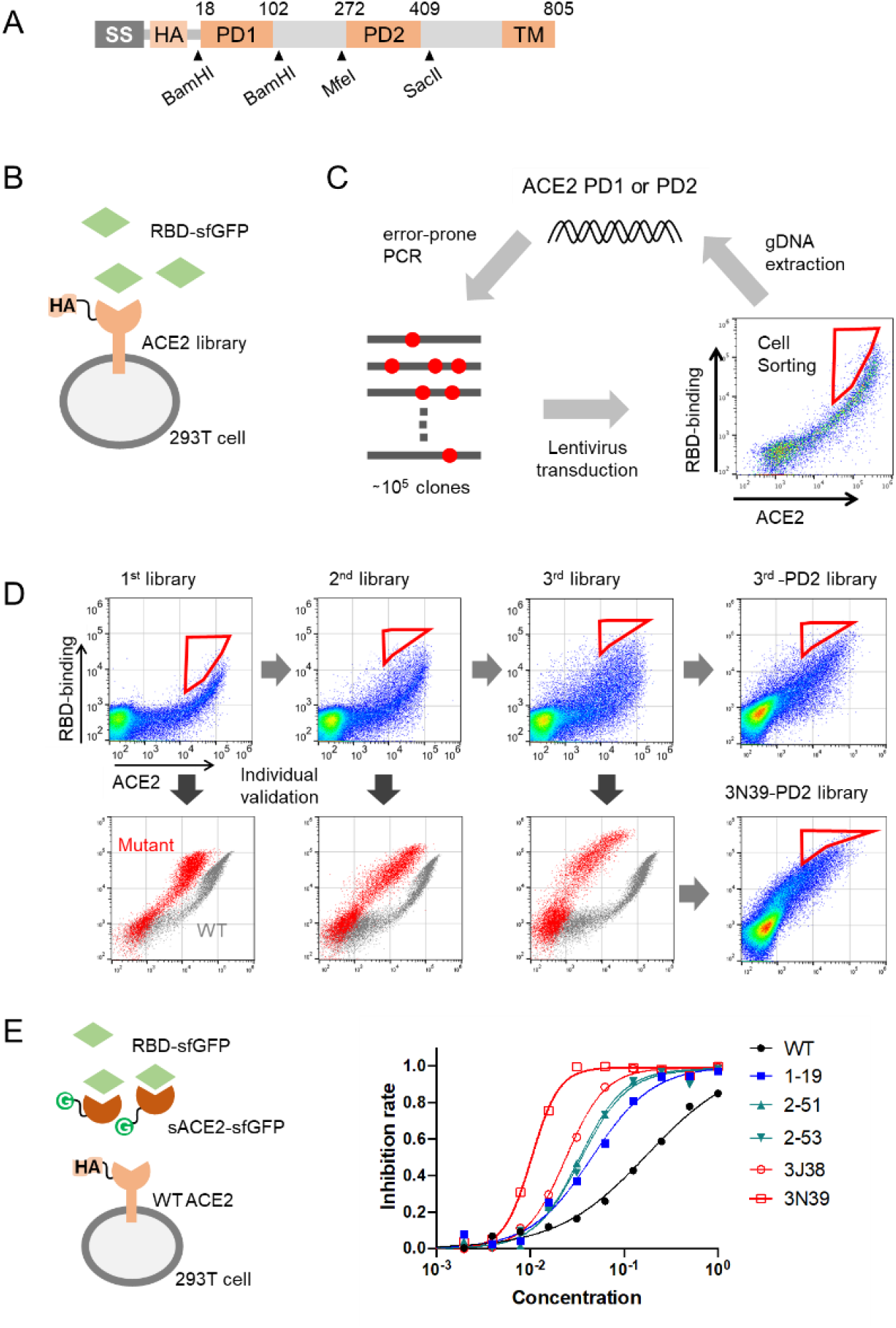
Directed evolution to generate high affinity ACE2 in 293T cells. (**A**) Full length ACE2 was optimized to fit screening. Synthetic signal sequence and HA tag were fused to mature ACE2 and restriction sites were introduced by optimizing codon optimization for the mutated fragment replacement. (**B**) ACE2 mutant library was expressed in 293T cells and incubated with the RBD of SARS-CoV-2 fused to superfolder GFP (sfGFP). (**C**) Error-prone PCR amplification of ACE2 protease domain induced random mutations. Mutant library-transduced cells were incubated with the RBD-sfGFP. Top 0.05 % population with high level of bound RBD-sfGFP was sorted and underwent DNA extraction, followed by next cycle mutagenesis. (**D**) PD1 mutagenesis and high affinity selection were performed 3 times and followed by PD2 random mutation. Top 0.05% population were harvested and reconstructed into the backbone plasmid to be verified individually. (**E**) Neutralization activity against the RBD was evaluated with sACE2-sfGFP and the RBD-sfGFP. Serial dilution of sACE2-sfGFP was analyzed with the 50-fold diluted RBD-sfGFP in flow cytometry. The experiments were independently performed twice and similar results were obtained. One representative data were shown.

Random mutagenesis screening for PD1 was performed 3 times and mutated sequences from top 0.05% population were reconstructed into the backbone plasmid, and expressed in 293T cells individually. The binding capacity to the RBD-sfGFP was verified in 100-300 clones. As the selection cycle advances, the two-dimensional distribution of library cells in flow cytometry became broader and higher in RBD-binding signal, and individual clone validation identified several mutants with higher binding capacity (Fig.1D). To evaluate the neutralization activity in the form of sACE2, we first generated fusion protein of the soluble extracellular domain of mutagenized ACE2 residues 18-614 and sfGFP (sACE2-sfGFP) and used them to compete with the cell-surface WT ACE2 for the RBD binding. To this end, the concentration of each mutant sACE2-sfGFP in the cultured medium from transfected cells was quantitatively standardized with sfGFP signal, serially diluted, preincubated with RBD-sfGFP for 30 min, and then transferred to WT ACE2 expressing 293T cells. After 30 min, the RBD-bound cells were analyzed in flow cytometry. Higher neutralization activity against the RBD was confirmed according to the accumulation of mutagenesis cycles (Fig. 1E, table S1).

Second mutagenesis based on the top hit of the first screening, 1-19 mutant, was also performed, but the distribution of the library cells expanded little, and we could not isolate clones showing higher binding signals than the bulk of top 0.05% (fig. S1). We next performed PD2 mutagenesis in both the bulk of top 0.05% and one of the highest mutants of the 3^rd^ library, the clone 3N39. Again, the binding distribution of the PD2 library cells was similar to the basal cells (Fig.1D), suggesting the inability of this strategy to obtain clones with higher binding activity to RBD. A recent study reported, via deep mutational scanning, that several specific mutations in PD2 were enriched in high RBD-binding clones (*16*). Even when we applied these mutations in 3N39, it failed to further improve the capacity of the RBD neutralization (fig. S2). In the end, we identified 3 highest neutralizing mutants, 3N39, 3J113 and 3J320 in the top population from 3rd library. To identify essential mutations, each mutation was altered to WT in these 3 mutants individually or in combination (figs. S3). Mutants composed of essential mutations were referred to as v2 (Fig. 2A) and we characterized their binding affinities by surface plasmon resonance (SPR). The K_D_ value of WT sACE2 was 17.63 nM, whereas those of mutants were determined to be from 0.29 nM to 3.98 nM (Fig. 2B), confirming that the increased RBD-binding activity of these mutants were in fact due to the increased binding affinity up to ∼100-fold. The SPR sensorgrams indicates that high affinity mutants all have very slow off-rate compared to the WT ACE2. It is commonly believed that ACE2 binding can occur when RBD is in a “up” conformation in the spike trimer, and most of the cryo-EM images of the ACE2-spike complex have only one ACE2 bound per spike trimer (*19*). In fact, a mass photometry analysis of WT ACE2-spike trimer complex reveals that it exists as the mixture of complexes with varying ratios, with 1:1 complex as the major species. In sharp contrast, spike trimer incubated with either 3N39 or 3N39v2 mutants behaved predominantly as 3:1 stoichiometric complex (fig. S4), indicating that the mutants can saturate all three RBDs on the spike trimer due to their slow dissociation.

**Fig 2.**
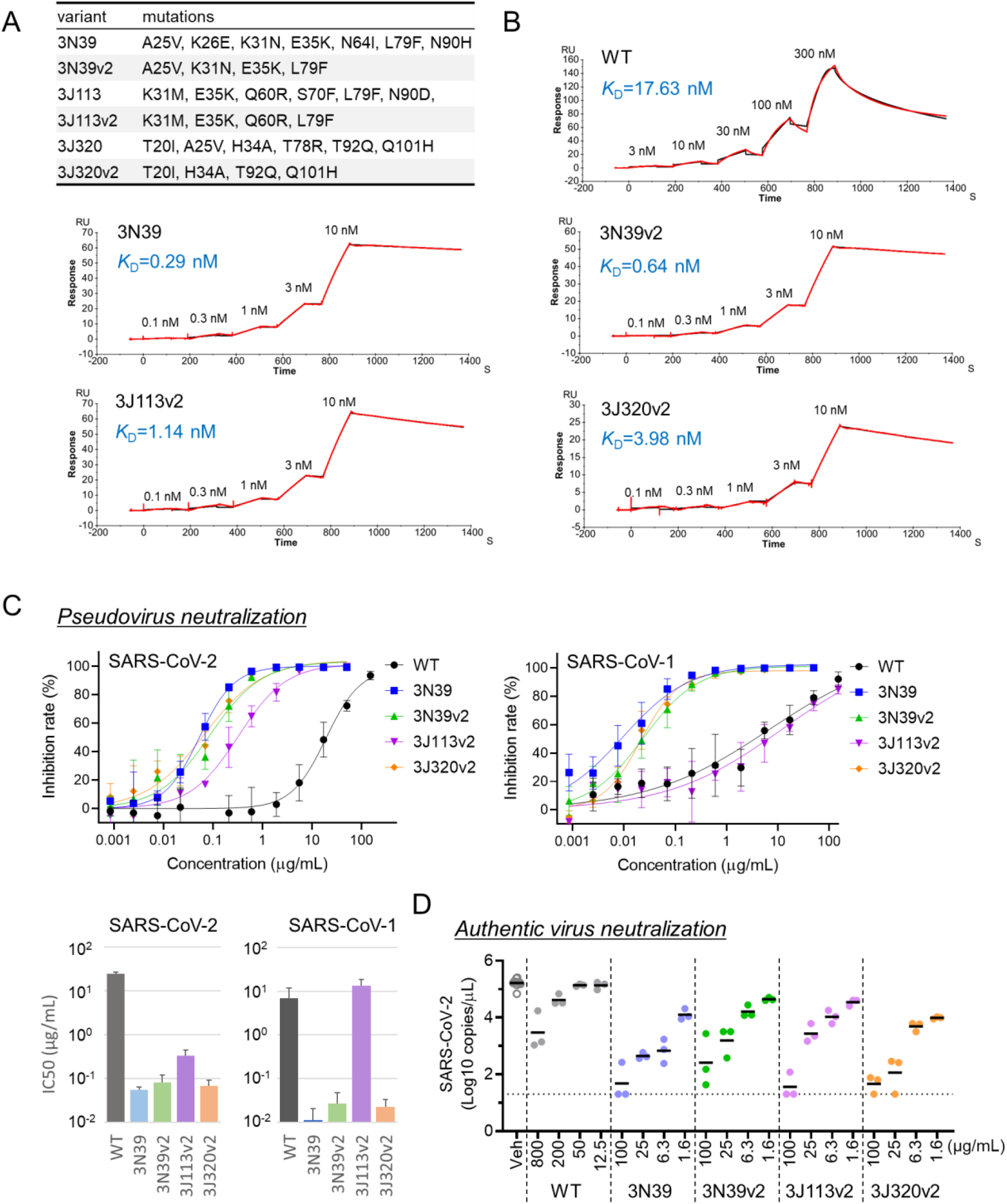
High affinity modified sACE2s have potential to neutralize SARS-CoV-2. (**A**) Table of all ACE2 variants with mutations. (**B**) Kinetic analysis of sACE2-His binding to RBD-Fc was analyzed by surface plasmon resonance (SPR). (**C**) Neutralization potency of sACE2-Fc against SARS-CoV-2 or SARS-CoV-1 pseudotyped lentivirus was measured in 293T/ACE2 cells. Data are mean ± SD of n = 3 biological replicates. (**D**) Neutralization potency to authentic SARS-CoV-2 was analyzed in Vero6E/TMPRSS2 cells. Data are mean of n = 3 technical replicates.

Having a few high affinity ACE2 mutants in hand, we next decided to assess their potentials as anti-SARS-CoV-2 therapeutic agents. First, we confirmed that none of the mutants were structurally unstable compared with WT ACE2 by measuring the Tm values in a thermal denaturation experiment (fig. S5). In fact, mutants 3J113v2 and 3J320v2 showed Tm values higher than the WT. Next we formulated our high affinity mutant sACE2s as human IgG1 Fc fusion (ACE2-Fc), as recombinant soluble ACE2 (rsACE2) monomer was reported to have a fast clearance rate in human blood with a half-life of hours (*20, 21*). We then evaluated their efficacy in neutralizing SARS-CoV-2 infections. First, we confirmed that mutant ACE2-Fcs directly inhibited the binding of full-length spike trimer to cell-surface ACE2 with more than 100-fold increase in the blocking efficacy from the WT (fig. S6). Pseudotyped SARS-CoV-2 neutralization assay in ACE2-expressing 293T cells exhibited that IC_50_ values of WT, 3N39, 3N39v2, 3J113v2 and 3J320v2 were 24.8, 0.056, 0.082, 0.33 and 0.068 *μ*g/ml, respectively (Fig. 2C, left). All mutants except for 3J113v2 were also capable of strongly inhibiting the infection of pseudotyped SARS-CoV-1, another ACE2-dependent coronavirus, indicating wide-range of therapeutic potential of 3N39v2 and 3J320v2 mutants (Fig. 2C, right). When the neutralization potential against the authentic SARS-CoV-2 in TMPRSS2-expressing VeroE6 cells was evaluated, each mutant ACE2-Fc demonstrated significant neutralizing effect even in 100-fold lower concentration than WT (Fig. 2D).

In order to know the structural basis for the affinity enhancement, we next determined the structure of the highest affinity ACE2 mutant, 3N39, in complex with the SARS-CoV-2 RBD. The crystal structure of the complex was solved at 3.2-Å resolution (Fig. 3A). Two 3N39-RBD complexes are contained in the asymmetric unit and they are structurally indistinguishable with a root mean square deviation (RMSD) value of 0.176 Å for 672 Cα atoms. The crystal structure confirms that 3N39 binds to RBD using the same interface as WT ACE2 reported previously (*15, 22-24*). Among the 7 mutated residues in 3N39, the side chains of E26, I64, and H90 are exposed to solvent and not involved in either inter- or intra-molecular interactions, corroborating their nonessential nature in the affinity enhancement. In contrast, mutation sites K31N and E35K are located at the center of the binding interface. In the WT structure, E35 forms an intramolecular salt bridge with the K31 in the preceding helical turn (Fig. 3B, right), making it less favorable for forming direct intermolecular hydrogen bonding with Q493 in the RBD. This salt bridge is lost by the simultaneous mutations of K31N and E35K in 3N39, and K35 now forms a direct inter-molecular hydrogen bond exclusively with Q493 of RBD (Fig. 3B, left). This results in small but significant ∼1 Å approach of the α1 helix of ACE2 toward RBD, likely contributing to the affinity enhancement of the mutant. As for L79F and A25V, both mutations are conservative in nature but accompany acquisition of larger sidechains. In WT ACE2, L79 forms a small hydrophobic pocket with Y83 and M82 that accommodates F486 of RBD, which has been suggested to be a key determinant of SARS-CoV-2 that gives higher affinity than SARS-CoV-1 (Fig.3C, right) (*22*). This hydrophobic contact became even more extensive due to the L79F mutation in 3N39. In addition, A25V mutation fills a space at the back of the pocket together with L97 (Fig.3C, left). It is thus assumed that these mutations collectively lead to the 100-fold increase in the overall affinity, which is equivalent to 2.76 kcal/mol.

**Fig 3.**
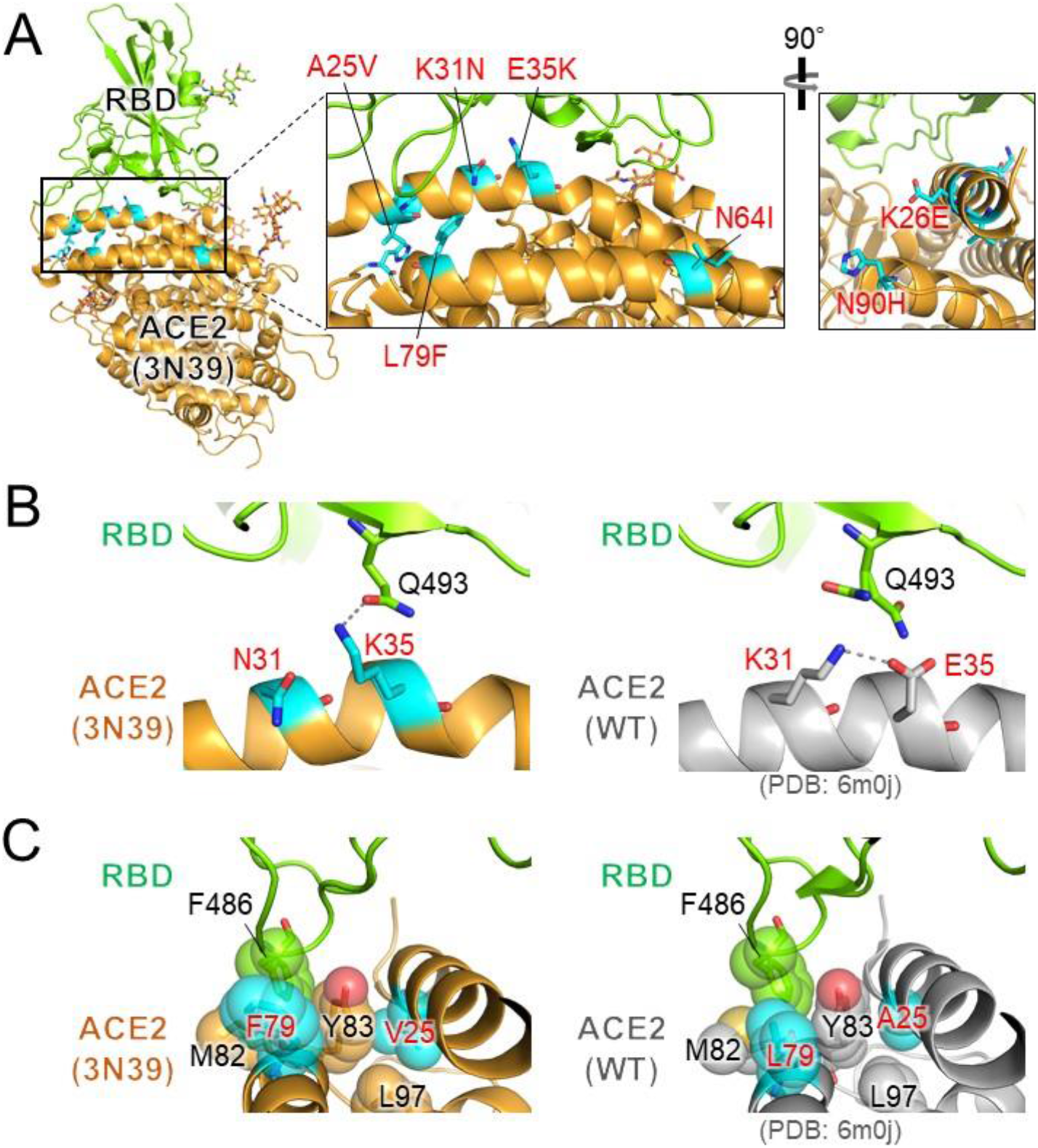
Structural analysis of 3N39 mutant in complex with RBD. (**A**) Overall structure. 3N39 ACE2 and RBD are shown in orange and green, respectively. The mutated residues in 3N39 are shown as cyan stick models. The expanded views of the PD1 region are provided in the inset. (**B**) Structural comparison of the K31N/E35K mutation site in 3N39 (left panel) with its corresponding site in WT (right panel). Hydrogen-bonding interactions (within 3.0 Å) are indicated by dashed lines. (**C**) Structural comparison of the L79F/A25V mutation site in 3N39 (left panel) with its corresponding site in WT (right panel). F486 residue of RBD and hydrophobic residues composing the F486 binding pocket of ACE2 are shown as stick models with transparent sphere models.

In the crystal structure, 3N39 adopts a “closed” conformation where the enzymatic active site is completely inaccessible due to the closure of the substrate binding groove (fig. S7A, left). This is very different from the “open” structures of WT ACE2 found in both RBD-bound (e.g., 6m0j) and unbound (e.g., 1r42) states (fig. S7A, right). Interestingly, the same closed structure was reported with WT ACE2 bound by an inhibitor MLN-4760 (Fig. S7B, center)(*25*). In fact, superposition of the two structures at the protease domain resulted in a surprisingly low RMSD value of 0.471 Å for 540 Cα atoms (Fig. S7B). In the 3N39 ACE2 structure, the active site Zn is intact but there was no density corresponding to any substrate-like compounds in the pocket except for a sulfate ion (fig. S7C). This observation made us to wonder if the closed conformation of the 3N39 mutant contributed to its increased RBD binding ability, although the active site groove was far away from the RBD-binding interface (fig. S7A). In order to test this idea, we designed an ACE2 mutant that is fixed in the closed conformation by introducing S128C/V343C double mutations in WT ACE2 (fig. S7D). Successful formation of the designed disulfide bond was inferred by an upward band shift of the mutant in a non-reducing SDS-PAGE (fig. S7E), as well as its complete loss of ACE2 activity (fig. S7F). When this mutant was evaluated for its RBD-binding neutralization activity, it showed identical dose response curve with the WT ACE2 (fig. S7G), indicating that the closed conformation *per se* is not responsible for the enhanced affinity. However, this observation gives us an important lesson that the inhibitor-blocked ACE2 in the closed conformation can still engage SARS-CoV-2 RBD, suggesting that caution must be taken when considering the possibility of using ACE2 inhibitors as the therapeutics against coronavirus infection (*26*).

There is a general concern in antiviral therapies that pathogenic virus could acquire drug resistance due to the frequent mutation. Actually, it was reported that SARS-CoV-2 escape mutation arose rapidly during the culture with a neutralizing antibody(*9*, 10). We evaluated the occurrence of escape mutation using authentic SARS-CoV-2 virus. At the first passage, 0.1 MOI virus was added to the culture in the presence of serially diluted mutant ACE2-Fc or recombinant monoclonal antibody (clone H4) isolated from a convalescent patient (*27*), and a total of 3 × 10^5^ copies of amplified virus from partially neutralized well was transferred to the next passage (Fig. 4A). Consistent with previous report (*9*, 10), the virus pool after the treatment with single antibody became insensitive to the highest concentration of the same antibody at the fourth passage, suggesting the acquisition of the antibody resistance. In contrast, the neutralizing capacity of the mutant ACE2-Fcs remained very high against the passaged virus pool, indicating the lack of emergence of escape mutations (Fig. 4B).

**Fig 4.**
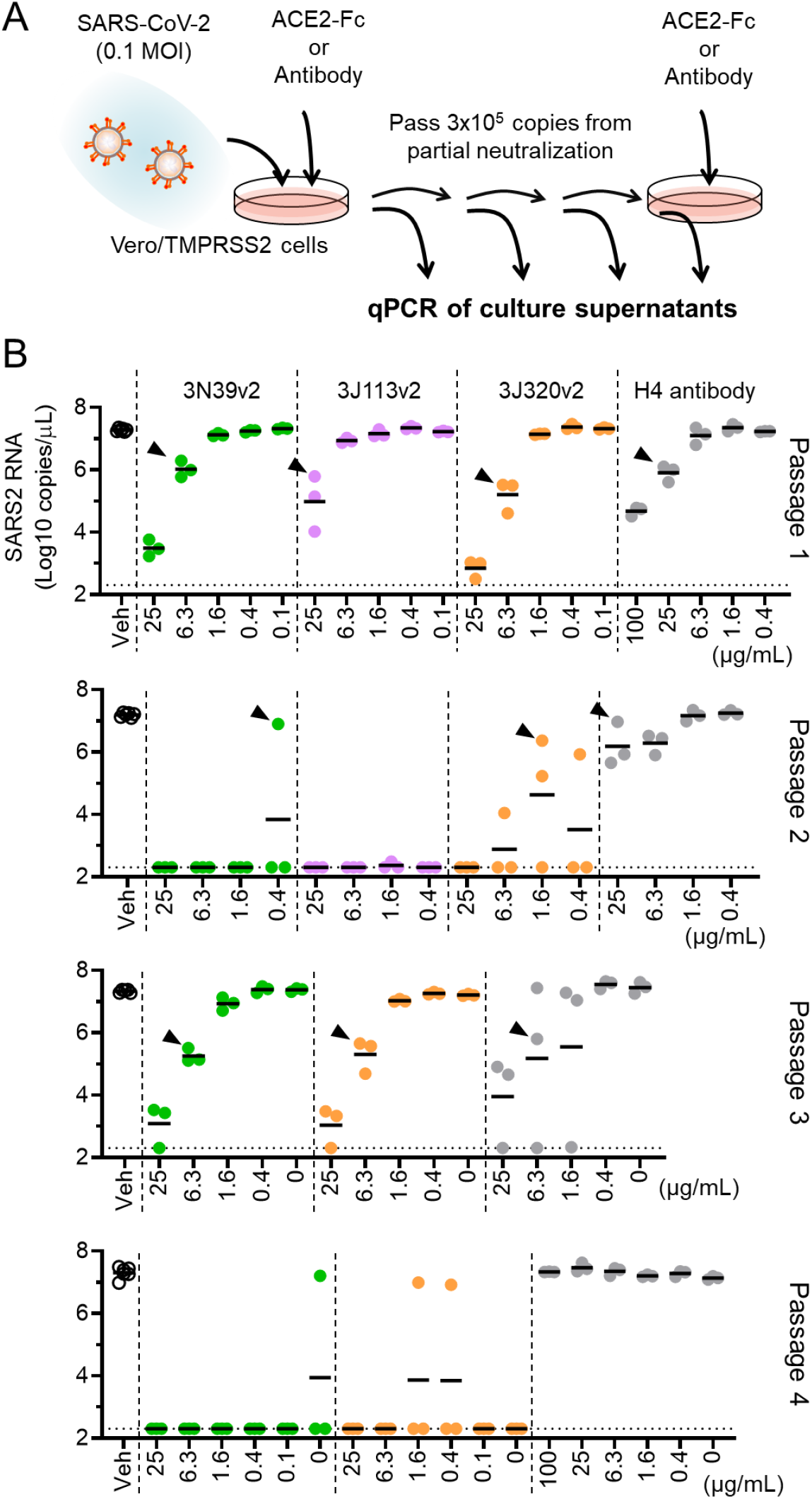
No emergence of escape mutation under the treatment of modified ACE2-Fc. (**A**) Protocol of generating escape mutation in SARS-CoV-2. At first, 0.1 MOI of SARS-CoV-2 was cultured in Vero6E/TMPRSS2 cells with indicated concentration of ACE2-Fc or H4 antibody, then a total of 3 × 10^5^ copies of virus in partially neutralized well was passed in the presence of ACE2-Fc or H4 antibody dilution. Supernatants were collected from each well and analyzed virus RNA copy number by quantitative PCR. (**B**) The copy number of SARS-CoV-2 genome RNA in cultured medium was analyzed in each passage. The virus from indicated well (arrow head) was passed and escape mutant expansion was observed only in H4 antibody at passage 4. The test for 3J113v2 was discontinued due to no growth of virus at passage 2.

Among three v2 mutant ACE2-Fcs showing similar neutralizing capacity and structural stability, we selected 3N39v2-Fc based on its high expression efficiency (fig. S8) to be used in the evaluation of therapeutic potential in hamster model of COVID-19 characterized by rapid weight loss and severe lung pathology (*28*). Since pharmacokinetic analysis of ACE2-Fc exhibited short half-life despite the effective distribution to lung tissues (figs. S9), 3N39v2-Fc or control-Fc were intraperitoneally administered 2 h after intranasal 1.0×10^6^ plaque forming units (PFU) virus challenge as a therapeutic application (Fig. 5A). Hamsters that received control-Fc lost 4.3% of body weight, whereas those treated with 3N39v2-Fc gained 7.3% similarly to the unchallenged control 5 days post-infection (Fig. 5B). In micro-CT before necropsy, the control group showed multi-lobular ground glass opacity mainly in the cranial portion, conversely, lung abnormalities were limited in the treated one (Fig. 5C, movies S1-S3). The content of SARS-CoV-2 in lungs was evaluated as functional virus particle and genome RNA copy number and the both were significantly decreased in the treatment with 3N39v2-Fc (Fig. 5D). We also performed histopathological analyses of infected hamsters. The control hamsters showed severe interstitial pneumonia characterized by widespread infiltration of inflammatory cells, alveolar septal thickening, and alveolar hemorrhage, whereas 3N39v2-Fc treatment obviously reduced lung pathologies and viral antigen (Figs. 5E and F). Consistent with these results, the expression of inflammatory or chemotactic cytokines was attenuated in the treated group (Fig. 5G).

**Fig 5.**
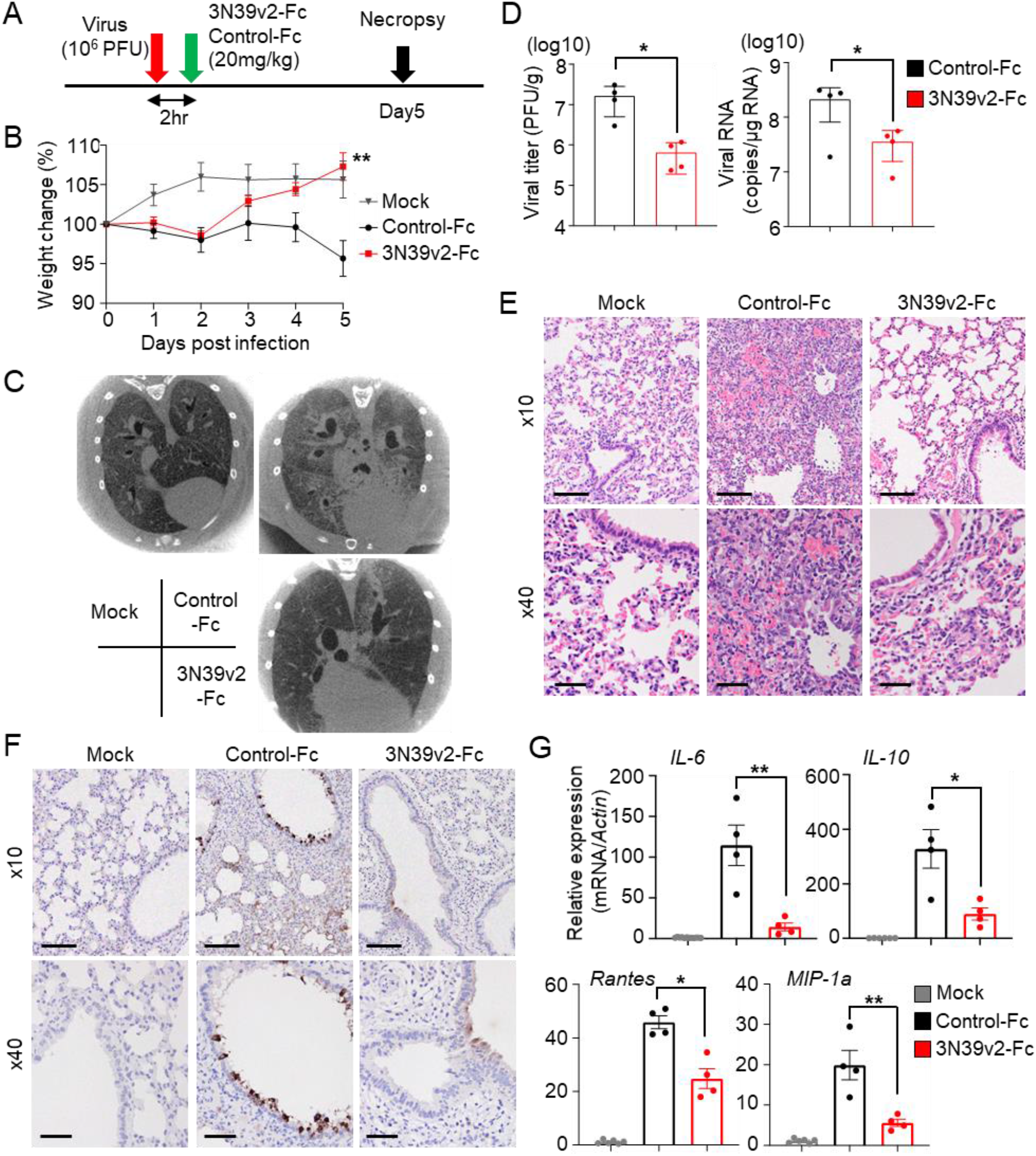
Therapeutic efficiency of 3N39v2-Fc in a COVID-19 Hamster Model. (**A**) Schematic overview of the animal experiment. (**B**) Percent body weight change was calculated from day 0 for all hamsters. Data are mean ± SEM of n = 4. (**C**) Axial CT images of the thorax 5 days after infection. (**D**) Quantification of plaque-forming units (PFU) from lung homogenates and genomic SARS-CoV-2 RNA as copies per *μ*g of cellular transcripts. (**E**) H&E staining and (**F**) SARS-CoV-2 antigen staining of hamster lung lobes. Scale bars, 100 *μ*m (upper panel) and 40 *μ*m (lower panel). (**G**) mRNA expression of inflammatory or chemotactic cytokines in hamster lung lobes. Significance levels are shown as *p < 0.05, **p < 0.01,

In addition to the decoy receptor, ACE2-Fc might have a protective effect on acute lung injury through proteolysis of angiotensin II (*12*, 29), but could have off-target vasodilation effect in the high-dose administration. Our ACE2 mutants exhibited similar catalytic activity with WT (fig. S10) and the closed-type mutation in the binding pocket of ACE2 exhibited no enzymatic activity (fig. S7F) and intact neutralizing activity in both WT and 3N39v2 background (fig. S7G). Whether catalytic function should be active or deleted remains to be investigated.

High affinity modified ACE2 fused with Fc is the promising strategy to neutralize SARS-CoV-2. The time frame for running one cycle of mutagenesis and sorting was just one week in our system, and we succeeded in developing optimized mutants in a couple of months independently of patients-derived cells or tissues. Thus, our system can rapidly generate therapeutic candidates against various viral diseases and may be well suited for fighting against future viral pandemics.

## ACKNOWLEDGEMENTS

We would like to thank Sho Hashimoto, Toshiyuki Nishiji, Tomohiro Hino, and Keiko Tamura-Kawakami for the construction of expression vectors; Yukiko Takemura for production of ACE2 proteins; Tomoya Kitani and Daisuke Ueno for the validation experiment; Shunta Taminishi for NGS sample preparation; Takeshi Yaoi for helpful discussion and support; Takao Hashiguchi for kind gift of plasmids coding for SARS-CoV-1 and 2 Spike protein.

## Author contribution

A.H. designed the research; Y.H., N.I., and A.H. performed the directed evolution screening; Y.H. performed RBD neutralization assay and analyzed pharmacokinetics; N.I. performed pseudovirus neutralization assay and ACE2 catalytic activity assay; Y.K., D.M., and S.N. performed and analyzed next-generation sequencing; T.A., E.M., and J.T. purified and prepared the proteins; T.A. performed Biacore assays, Mass photometry, and X-ray structure analysis; E.M. conducted neutralization assay for S trimer; J.T. performed thermal shift assay; E.O. performed SARS-CoV-2 neutralization assay; T.S., Y.I, F.S., and T.O. conducted SARS-CoV-2 experiments in the cell culture and hamsters; Y.S. performed histological analysis; O.M., Y.M., S.M., T.O., J.T., and A.H. supervised the research; A.H. J.T., and T.O. wrote the manuscript; all authors discussed the results and commented on the manuscript.

## Data and materials availability

Crystallographic coordinates and structure factors for 3N39-RBD complex have been deposited in the Protein Data Bank under accession code 7DMU.

## SUPPLEMENTARY MATERIALS

### Materials and Methods

Figs. S1 to S10

Tables S1 and S2

Movies S1 to S3

References *(30-40)*

## Supplementary Materials

### Materials and Methods

#### Cell culture

Lenti-X 293T cells were purchased from Clontech and cultured at 37 °C with 5% CO_2_ in Dulbecco’s modified Eagle’s medium (DMEM, WAKO) containing 10% fetal bovine serum (Gibco) and penicillin/streptomycin (100 U/ml, Invitrogen). VeroE6/TMPRSS2 cells were a gift from National Institutes of Biomedical Innovation, Health and Nutrition (Japan) and cultured at 37 °C with 5% CO_2_ in DMEM (WAKO) containing 5% fetal bovine serum (Gibco) and penicillin/streptomycin (100 U/ml, Invitrogen). All the cell lines were routinely tested negative for mycoplasma contamination.

#### Plasmids

The mature polypeptide (a.a. 18-805) of human ACE2 (GenBank NM_021804.1) was cloned into the KpnI-XhoI sites of pcDNA4TO (Invitrogen) or the XbaI-SalI sites of pLenti puro with a N-terminal synthetic leader (MWWRLWWLLLLLLLLWPMVWA)(*30*), HA-tag, and linker (GSGG). Some codon optimizations were introduced to generate BamHI and SacII and destroy original BamHI and SacII restriction sites. Soluble ACE2 (a.a. 1-615) fused to superfolder GFP (sfGFP) with a linker (GSGGSGSGGS) was placed between the BamHI-XhoI sites of the former plasmid. Equivalent sACE2 constructs were cloned with TEV protease cleavage site and 8 histidine tag or the Fc region of IgG1 into the KpnI-BstBI sites of pcDNA3.1-MycHis (Invitrogen) or the HindIII-BamHI sites of p3xFLAG-CMV-14 (Invitrogen). A codon-optimized RBD (a.a. 333-529) of SARS-CoV-2 Spike fused to sfGFP was obtained from addgene #141184 (*16*) and cloned into the KpnI-XhoI sites of pcDNA4TO (Invitrogen) with a N-terminal synthetic leader, HA-tag, and linker (GSGG).

#### Protein production and purification

sACE2-His, sACE2-Fc, and RBD-Fc were expressed using the Expi293F cell expression system (Thermo Fisher Scientific) according to the manufacturer’s protocol. His-tagged and Fc-fused proteins were purified from conditioned media using the Ni-NTA agarose (QIAGEN) and the rProtein A Sepharose Fast Flow (Cytiva), respectively. Fractions containing target proteins were pooled and dialyzed against phosphate buffered saline (PBS). The full-length spike timer protein was also produced in Expi293F cells by stably transfecting expression vector coding for the entire ectodomain portion of SARS-CoV-2 spike protein (residues 1-1212 with stabilizing mutations R684G, R685S, R678G, K998P, and V999P) with a C-terminal fibritin trimerization motif and hexahistidine tag (a gift from Takao Hashiguchi, Kyushu University). The spike trimer was purified from the culture supernatants by using Ni-NTA agarose followed by dialysis against PBS. An anti-RBD monoclonal antibody (clone H4) isolated from a convalescent patient (*27*) were formulated in the form of human IgG1/kappa by using synthetic DNA coding for the variable regions of heavy and light chains taken from the publicly available amino acid sequences, and recombinantly produced in Expi293F cells as above.

#### Library preparation

Error-prone PCR was performed in ACE2 residues 18-102 and 272-409 independently using GeneMorph II Random Mutagenesis Kit (Agilent). A 0.1 ng of plasmid was used as a PCR template and generated mutations with an average of about one mutation per 100 bp in 35-cycle reaction. The plenti ACE2 vector was digested with BamBI or MfeI-SacII (NEB) for ACE2 residues 18-102 or 272-409, respectively with alkaline phosphatase (Fermentas) at 37 °C for 2 hr and gel-purified on a Gel and PCR Clean-up kit (TAKARA). A 160 *μ*l NEBuilder ligation reaction (NEB) was performed using 5 ng of the gel-purified inserts and 10 ng of the vector, then purified on a Gel and PCR Clean-up kit (TAKARA) and eluted by a 20*μ*l of distilled water. From the ligation, 400 *μ*g (∼8*μ*l) of the purified reaction was transformed into 100 ul of electrocompetent cells (Lucigen) and expanded according to the manufacturer’s protocol with 1500V electroporation by ECM 399 (BTX). A 1,000-fold dilution of the full transformation was plated to estimate the scale of mutant library.

#### Virus production

Ten-centimetre plates of 70% confluent Lenti-X 293T (Clontech) cells were transfected with 9 *μ*g of the plasmid library, 6 *μ*g of psPAX2 vector and 3 *μ*g pMD2.G using Fugene HD (Promega) according to the manufacturer’s instructions. Supernatant was collected after 48 h and then spun for 10 min at 4 °C at 3,000 rpm and then filtered with a 0.45 *μ*m low protein-binding filter (SFCA), and frozen at –80 °C. The titre of the virus was determined by using 293T cells followed by puromycin selection.

#### Library screening

The mutant library was transduced into 293T cells via spinfection. To find optimal virus volumes for achieving a multiplicity of infection (MOI) of 0.1–0.3, each library virus was tested by spinfecting 5×10^5^ cells with several different volumes of virus in the 6-well plate. Each well received a different titrated virus amount (usually between 50 and 500 *μ*l) along with a no-transduction control and infected cell rate was determined by anti-HA Alexa 594. Then, four 6-well plates, 1.2 × 10^7^ cells were centrifuged at 1,000 g for 1.5 hr at 37 °C.

Cells were sorted using a SH800 (SONY) 24 hr after spinfection. Approximately 5×10^7^ induced library cells were resuspended with complete medium and incubated for 30 minutes at 4°C with a 1/40∼1/160 dilution of medium containing RBD-sfGFP and a 1/4000 dilution of anti-HA Alexa 647 (clone TANA2, MBL). Cells were directly sorted on SH800 (SONY). The top 0.5 % of cells were sorted and their genomic DNA was extracted by NucleoSpin Tissue (TAKARA). Mutated ACE2 fragment was cloned into pcDNA4TO ACE2 for individual validation and also introduced random mutations again with error-prone PCR for further screening.

#### Flow Cytometry Analysis

The pcDNA4TO HA-ACE plasmid was transfected into 293T cells (500 ng DNA per ml of culture at 5 × 10^5^ / ml) using Fugene HA (Promega). Cells were analyzed by flow cytometry 24 hr post-transfection. To analyze binding of RBD-sfGFP to full length HA-ACE2, cells were washed, trypsinized, and resuspended with complete medium, then incubated for 30 min at 4 °C with a 1/40∼1/160 dilution of medium containing RBD-sfGFP and a 1/4000 dilution of anti-HA Alexa 594 (clone TANA2, MBL). Cells were directly analyzed on Attune NxT Flow Cytometer (Invitrogen).

To analyze binding of sACE2-sfGFP to RBD-sfGFP, a serial dilution of medium containing sACE2-sfGFP and a 1/40 dilution medium containing RBD-sfGFP were mixed and incubated for 1 hr at 4 °C, then HA-ACE2 expressing 293T cells were resuspended and incubated with the mixture and a 1/1000 dilution of anti-HA Alexa 594 (clone TANA2, MBL) for 30 min at 4 °C. Cells were directly analyzed on Attune NxT Flow Cytometer (Invitrogen).

Neutralizing activity of ACE2-Fc against soluble spike trimer binding to cell-surface ACE2 was evaluated as follows. First, varying concentrations (0.3 - 30 nM) of purified spike trimer was preincubated with 60 *μ*g/ml (∼315 nM) ACE2-Fc proteins at room temperature for 2 hr, followed by addition to Expi293F cells transiently transfected with human full-length ACE2 2 days before the experiment. After incubation on ice for 1.5 hr, the cells were washed 3 times with PBS and further incubated with Alexa Fluor 488-labeled anti-His tag antibody (MBL, catalogue # D291-A48) at 2 *μ*g/ml to stain the bound spike protein. Cells were analyzed on an EC800 system (Sony) and the data were processed with FlowJo (FlowJo, LLC).

#### Kinetic binding measurement using Biacore (SPR)

The binding kinetics of sACE2 (wild-type or mutants) to RBD were analyzed by SPR using a Biacore T200 instrument (Cytiva) in a single-cycle kinetics mode. Anti-human IgG (Fc) antibody was immobilized onto a CM5 sensor chip (Cytiva) using the Human Antibody Capture Kit (Cytiva) according to the method provided by the manufacturer. The RBD-Fc was captured on the measurement cell via the antibody at a density of ∼450 RU, while human IgG1-Fc was captured on the reference cell at a density of ∼200 RU. The binding was evaluated by injecting various concentrations of sACE2-His solutions in series using PBS containing 0.05% Tween 20 as a running buffer. The runs were conducted at 25 °C employing the following parameters; flow rate of 30 *μ*l/min, contact time of 120 sec, and dissociation time of 480 sec. After each run, the surface was regenerated by injecting the Regeneration solution contained in the kit for 30 sec. The binding curves of the measurement cell were subtracted with those of the reference cell, and used to derive kinetic binding values. The results were evaluated by using Biacore T200 evaluation software version 4.1.

#### Thermal shift assay

The thermal stability of the mutant ACE2 ectodomain proteins were evaluated by differential scanning fluorimetry as follows. Purified ACE2-His proteins were diluted to 200 *μ*g/ml in PBS and placed in 0.2-mL white PCR tubes (Bio-Rad, TLS0851) at 20 *μ*l/tube. After adding 1*μ*l/tube of SYPRO™ Orange protein gel stain solution (Invitrogen, S6651) diluted with water at 1:150, the tubes were placed in a Bio-Rad CFX96 thermal cycler Real-Time Detection System. Thermal denaturation curves from 25 °C to 95 °C (ramp rate of 1.27 °C/min at 0.5-°C intervals with an equilibration of 5 sec at each temperature before measurement) were acquired by measuring fluorescence intensities using the FRET channel with excitation from 450 to 490 nm and detection from 560 to 580 nm. All data were exported and plotted in Microsoft Excel and the first derivative approach was used to calculate T_m_.

#### MassPhotometry

Binding stoichiometry between ACE2-His and spike trimer was determined by mass photometry (*31*). To this end, either WT or 3N39v2 mutant ACE2-His (∼75 kDa polypeptide with 5 N-glycans) was incubated with purified SARS-CoV-2 spike trimer (∼450 kDa polypeptide with 63 N-glycans) at 1: 1.2 molar ratio in PBS and placed on a microscope coverslip. MP data were acquired and analyzed using a One^MP^ mass photometer (Refeyn Ltd, Oxford, UK).

#### Crystallization

For crystallization, hingeless Fc was appended to RBD (RBD-noHg-Fc), and the protein was expressed in Expi293F cells in the presence of 5 µM *α*-mannosidase inhibitor, kifunensine (Cayman Chemical Co.). After purification using the rProtein A Sepharose, RBD-noHg-Fc was treated with His-tagged IdeS protease to cleave between the RBD and the Fc, followed by subjecting to the Ni-NTA agarose and the rProtein A Sepharose sequentially to remove His-IdeS and Fc, respectively, by recovering the unbound fractions. The sample was further purified by size-exclusion chromatography (SEC) on a Superdex 200 Increase 10/300 GL column (Cytiva) equilibrated with 20 mM Tris, 150 mM NaCl, pH 7.5. 3N39 mutant sACE2-His was also expressed in Expi293F cells in the presence of kifunensine. After Ni-NTA purification, the sample was purified by anion exchange chromatography on a MonoQ 5/50 GL column equilibrated with 20 mM Tris, pH 8.0. To obtain sACE2-RBD complex, the purified sACE2 and RBD were mixed at a molar ratio of 1 : 1.3 and subjected to SEC. Fractions containing the complex sample were concentrated to 9.2 mg/ml prior to crystallization. The diffraction quality crystal was grown under the condition of 1.6 M ammonium sulfate, 0.25 M lithium sulfate, and 0.05 M CAPS pH 10.5. The crystal was cryoprotected with the same buffer containing 25% ethylene glycol and used for data collection.

#### Data collection, phasing, and structure refinement

X-ray diffraction experiment was performed at beamline BL44XU of SPring-8 (Hyogo, Japan). Four datasets were collected at 100 K from a single crystal and were combined, processed, and scaled using the X-ray Detector Software (*32*). Initial phase was determined by molecular replacement method with PHASER v.2.8.1 (*33*) from the CCP4 package v.7.0 (*34*) using the crystal structure of RBD-sACE2(WT) (PDB: 6m0j) as a search model. The structural model was modified with COOT v.0.8.9 (*35*) and refined with PHENIX v.1.14 (*36*). Ramachandran plot analysis of the final structure with MOLPROBITY v.4.5 (*37*) showed that 96.5 and 3.3% of the residues are in favored and allowed regions, respectively. Data collection, processing, and refinement statistics are summarized in table S2. All structural figures were prepared with the PyMOL software v.2.3.4. (https://pymol.org/2/).

#### ACE2 Catalytic Activity Assay

Activity was measured using the Fluorometric ACE2 Activity Assay Kit (AnaSpec) with protein diluted in assay buffer to 100 ng/ml final concentration. Specific activity is reported as pmol MCA produced per 100 ng of enzyme. Fluorescence was read at 5 min interval on SpectraMax M2 (Molecular Devices).

#### Single-dose pharmacokinetics of ACE2-Fc

All animal experiments were performed according to procedures approved by Kyoto Prefectural University of Medicine Institutional Animal Care and Use Committees. A single 20mg/kg dose of ACE2-Fc was intraperitoneally injected to 8-week-old male C57BL/6JJcl mice (CLEA Japan) and whole blood was harvested from tail veil at indicated time points sequentially. Another cohort of mice was euthanized and whole blood and lungs were isolated 2 hr after injection. Heparinized whole blood was centrifuged, and plasma was collected and frozen. Lungs were perfused with 10 mL of cold PBS to remove blood before harvest, and then lung tissue lysates were prepared through mechanical disruption using Precellys tissue homogenizer (Bertin), followed by lysis with Cell Extraction Buffer PTR (Abcam). Lysates were cleared by centrifugation and were frozen for analysis. ACE2-Fc concentration was analyzed using human ACE2 ELISA Kit (Abcam; ab235649). Absorbance was read on Infinite F200 pro system (Tecan).

#### Pseudotyped virus neutralization assay

Pseudotyped reporter virus assays were conducted as previously described (*38*). A plasmid coding SARS-CoV-1 Spike was kindly gifted from Takao Hashiguchi (Kyushu University) and SARS-CoV-2 Spike was obtained from addgene #145032 (*15*), and deletion mutant CΔ19 (with 19 amino acids deleted from the C terminus) was cloned into pcDNA4TO (Invitrogen) to enhance virus titer(*39*). Spike-pseudovirus with a luciferase reporter gene was prepared by transfecting plasmids (CΔ19, psPAX2, and pLenti firefly) into LentiX-293T cells with Lipofectamine 3000 (Invitrogen). After 48 hr, supernatants were harvested, filtered with a 0.45 *μ*m low protein-binding filter (SFCA), and frozen at –80 °C. ACE2-expressing 293T cells were seeded at 10,000 cells per well in 96-well plate. Pseudovirus and three-fold dilution series of sACE2-Fc protein were incubated for 1 hr, then this mixture was administered to ACE2-expressing 293T cells. After 1 hr pre-incubation, medium was changed and cellular expression of luciferase reporter indicating viral infection was determined using ONE-Glo™ Luciferase Assay System (Promega) in 48 hr after infection. Luminescence was read on Infinite F200 pro system (Tecan).

#### Viruses

SARS-CoV-2 (2019-nCoV/Japan/TY/WK-521/2020) strain was isolated at National Institute of Infectious Diseases (NIID). SARS-CoV-2 were propagated in VeroE6/TMPRSS2 cells cultured at 37 °C with 5% CO_2_ in DMEM (WAKO) containing 10% fetal bovine serum (Gibco) and penicillin/streptomycin (100 U/ml, Invitrogen). The virus stock was generated by infecting VeroE6/TMPRSS2 cells at an MOI of 0.1 in DMEM containing 10% FBS; viral supernatant was harvested at two days post infection and the viral titer was determined by plaque assay.

#### SARS-CoV-2 neutralization assay

Vero-TMPRSS2 were seeded at 80,000 cells in 24 well plates and incubated for overnight. Then, SARS-CoV-2 was infected at MOI of 0.1 together with sACE2-Fc protein. After 2 hr, cells were washed by fresh medium and incubated with fresh medium for 22 hr. Culture supernatants were collected and performed qRT-PCR assay.

#### Syrian hamster model of SARS-CoV-2 infection

All animal experiments with SARS-CoV-2 were performed in biosafety level 3 (ABSL3) facilities at the Research Institute for Microbial Diseases, Osaka University. Animal Experimentation, and the study protocol was approved by the Institutional Committee of Laboratory Animal Experimentation of the Research Institute for Microbial Diseases, Osaka University (R02-08-0). All efforts were made during the study to minimize animal suffering and to reduce the number of animals used in the experiments. Four weeks-old male Syrian hamsters were purchased from SLC Japan. Syrian hamsters were anaesthetized by intraperitoneal administration of 0.75 mg kg-1 medetomidine (Meiji Seika), 2 mg kg-1 midazolam (Sandoz) and 2.5 mg kg-1 butorphanol tartrate (Meiji Seika) and challenged with 1.0 × 10^6^ PFU (in 60*μ*L) via intranasal routes. After 2 hr post infection, Control-Fc (20mg kg-1) or ACE2-Fc (3N39v2, 20mg kg-1) were dosed through intraperitoneal injection. Body weight was monitored daily for 5 days.

On 5 days post infection, all animals were euthanized and lungs were collected for histopathological examinations, virus titration and qRT-PCR. The structures of lungs were also observed by using micro computed tomography (*μ*CT)(ScanXmate-RB080SS110, Comscantecno Co.,Ltd.) with the following parameters: source voltage 80 kV; current, 0.1 mA; and voxel size, 0.1mm. The images of *μ*CT was reconstructed and visualized by ImageJ software.

#### SARS-CoV-2 virus plaque assays

VeroE6/TMPRSS2 cells were seeded on 24 well plates (80,000 cells/well) and incubated for overnight. The lung homogenates serially diluted by medium were inoculated and incubated for 2 hr. Culture medium was removed, fresh medium containing 1% methylcellulose (1.5mL) was added, and the culture was further incubated for 3 days. The cells were fixed with 4% Paraformaldehyde Phosphate Buffer Solution (Nacalai Tesque) and plaques were visualized by using a Crystal violet or Giemsa’s azur-eosin-methylene blue solution (Merck Millipore: 109204).

#### Quantitative RT-PCR

Total RNA of lung homogenates was isolated using ISOGENE II (NIPPON GENE). Real-time RT-PCR was performed with the Power SYBR Green RNA-to-CT 1-Step Kit (Applied Biosystems) using a AriaMx Real-Time PCR system (Agilent). The relative quantitation of target mRNA levels was performed by using the 2-ΔΔCT method. The values were normalized by those of the housekeeping gene, *β*-actin. The following primers were used: for *β*-actin; 5’-TTGCTGACAGGATGCAGAAG-3’ and 5’-GTACTTGCGCTCAGGAGGAG-3’, 2019-nCoV_N2 ; 5’-AAATTTTGGGGACCAGGAAC -3’and 5’-TGGCAGCTGTGTAGGTCAAC -3’, IL-6; 5’-GGA CAATGACTATGTGTTGTTAGAA -3’and 5’-AGGCAAATTTCCCAATTGTATCCAG -3’, MIP-1a; 5’-GGTCCAAGAGTACGTCGCTG -3’and 5’-GAGTTGTGGAGGTGGCAAGG -3’, Rantes; 5’-TCAGCTTGGTTTGGGAGCAA -3’and 5’-TGAAGTGCTGGTTTCTTGGGT -3’, IP-10; 5’-TACGTCGGCCTATGGCTACT -3’and 5’-TTGGGGACTCTTGTCACTGG -3’.

#### Quantitative RT-PCR of Viral RNA in the supernatant

The amount of RNA copies in the culture medium was determined using a qRT-PCR assay as previously described with slight modifications (*40*). In brief, 5 *μ*l of culture supernatants were mixed with 5 *μ*l of 2*×* RNA lysis buffer (2% Triton X-100, 50 mM KCl, 100 mM Tris-HCl [pH 7.4], 40% glycerol, 0.4 U/*μ*l of Superase•IN [Life Technologies]) and incubate at room temperature for 10 min, followed by addition of 90 *μ*l of RNase free water. 2.5 *μ*l of volume of the diluted samples was added to 17.5 *μ*l of a reaction mixture. Real-time RT-PCR was performed with the Power SYBR Green RNA-to-CT 1-Step Kit (Applied Biosystems) using a AriaMx Real-Time PCR system (Agilent).

#### Escape mutation study

VeroE6/TMPRSS2 cells were seeded on 96 well plates and infected with SARS-CoV-2 at MOI of 0.1 together with recombinant proteins. After 2 days post-infection, culture supernatants were collected, quantified copy numbers of viral RNA and a total of 3 x10^5^ copies of virus were further infected in naïve VeroE6/TMPRSS2 cells with recombinant proteins.

#### Hematoxylin and eosin staining and Immunohistochemistry

Lung tissues were fixed with 10% neutral buffered formalin and embedded in paraffin. 2 *μ*m tissue sections were prepared and stained with Hematoxylin and eosin (H&E). For immunohistochemical staining, 2 *μ*m thickness sections were immersed in citrate buffer (pH 6.0) and heated for 20 min with a pressure cooker. Endogenous peroxidase was inactivated by immersion in 3% H_2_O_2_ in PBS. After treatment with 5% BSA in PBS for 30 min at room temperature, the sections were incubated with mouse anti-Spike protein antibody (1:300, clone 1A9, Genetex, Irvine, CA, USA). For secondary antibody, EnVision+ system-HRP-labeled polymer anti-mouse or anti-rabbit (Dako, Carpinteria, CA, USA) were used. Positive signals were then visualized by peroxidase–diaminobenzidine reaction and sections were counterstained with hematoxylin.

**Fig S1.**
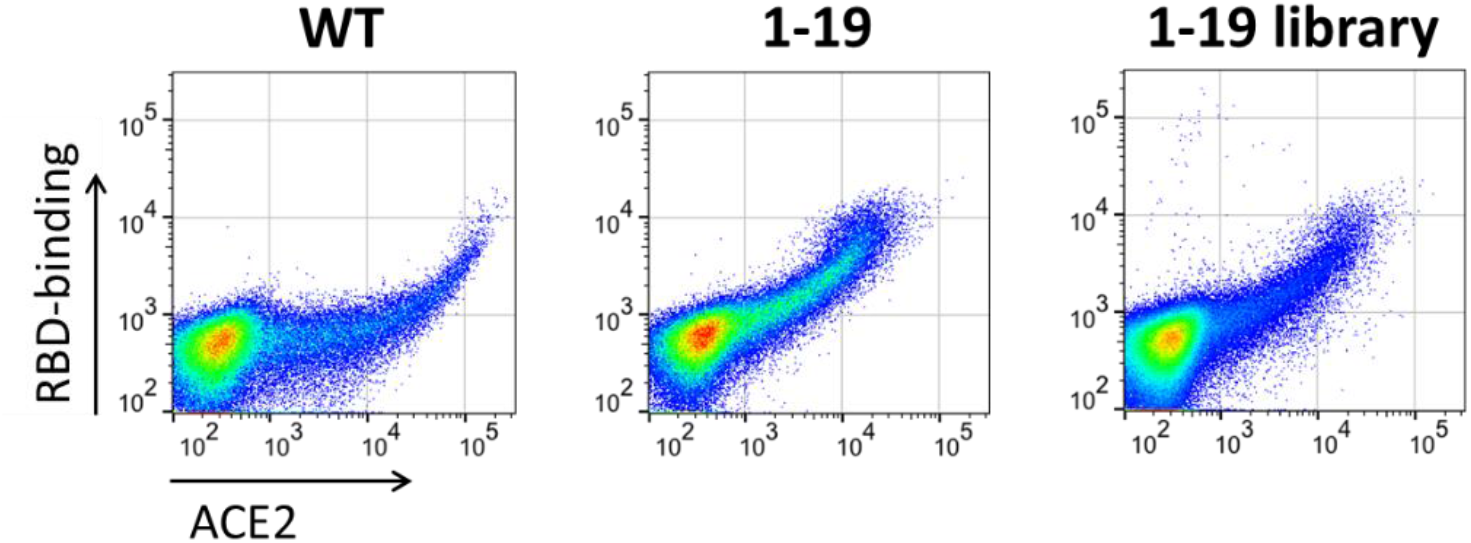
Second mutagenesis based on the top hit, 1-19 mutation in first screening. The distribution of mutagenesis library from 1-19 mutation was narrow and provided no better mutant than original 1-19 mutant in the individual validation.

**Fig S2.**
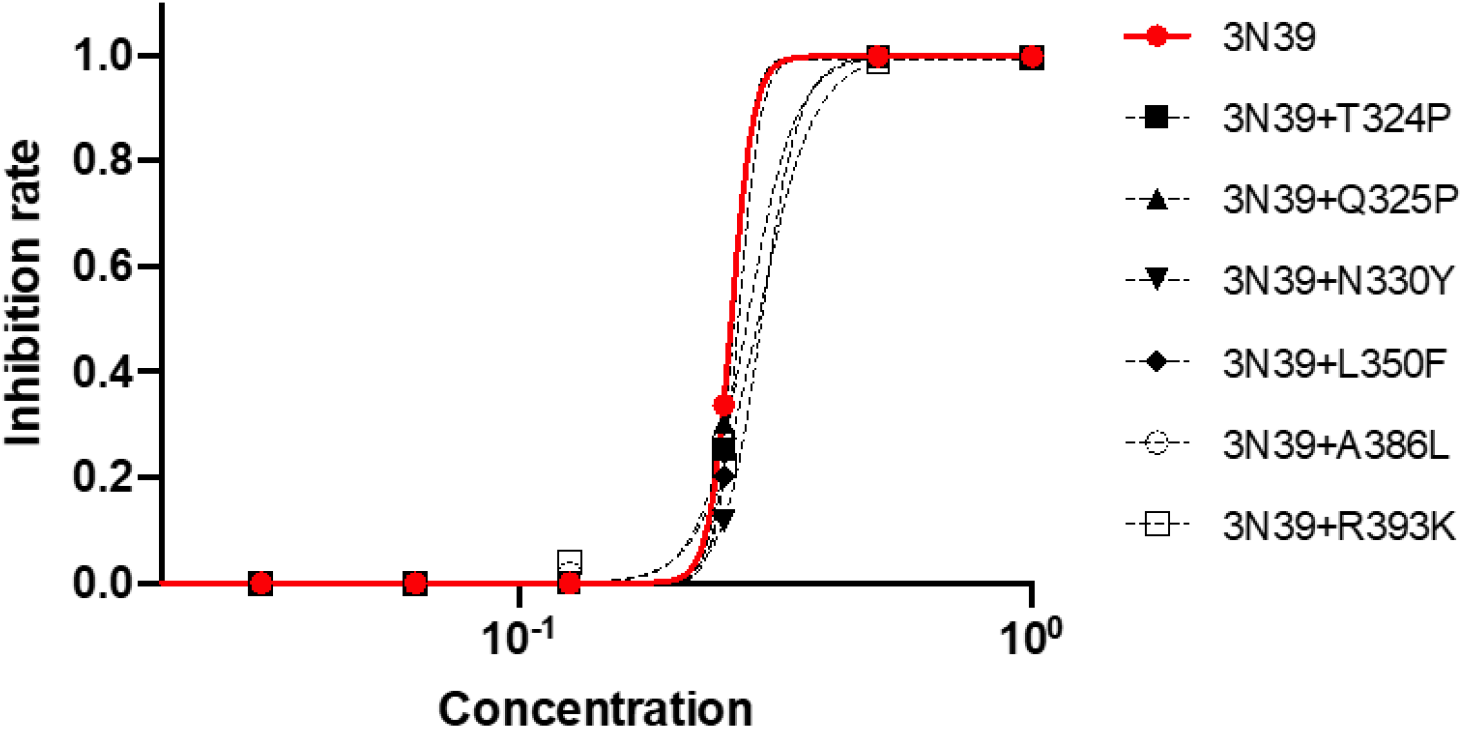
Additional PD2 mutation in 3N39 mutant. Induction of affinity-enhancing mutation in PD2 from Procko’s paper (*16*) failed to improve the capacity of the RBD neutralization. The experiments were independently performed twice and similar results were obtained. One representative data were shown.

**Fig S3.**
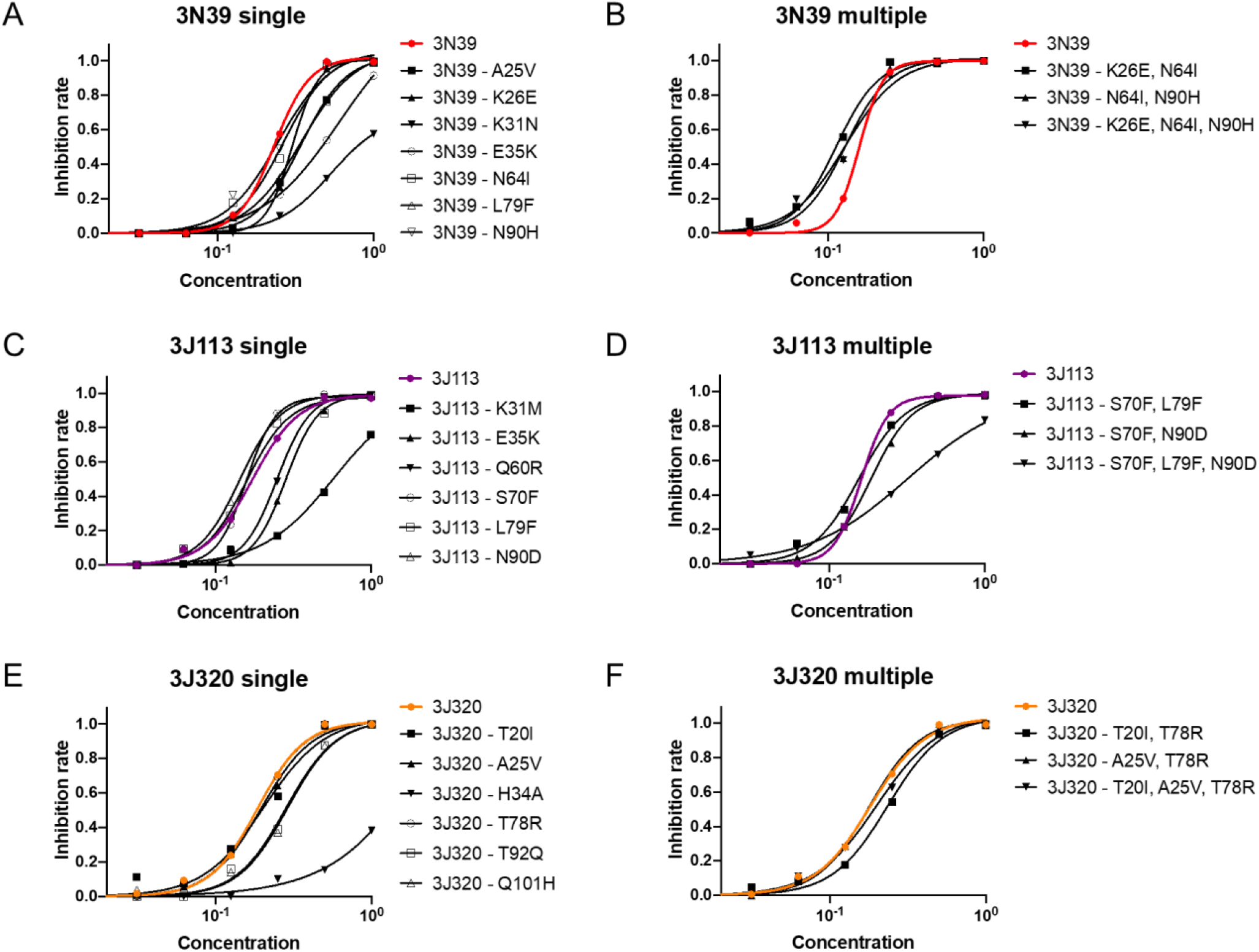
The contribution of each mutation in high affinity mutants. Serial dilution of sACE2-sfGFP was analyzed with 20-fold dilution of the RBD-sfGFP in flow cytometry.(**A-F**) Each mutation was recovered to wild-type and then analyzed the alteration of neutralization capacity individually in 3N39 (**A**), 3J113 (**C**), and 3J320 (**E**) and also in combination with non-essential mutations in 3N39 (**B**), 3J113 (**D**), and 3J320 (**F**). The experiments were independently performed twice and similar results were obtained. One representative data were shown.

**Fig S4.**
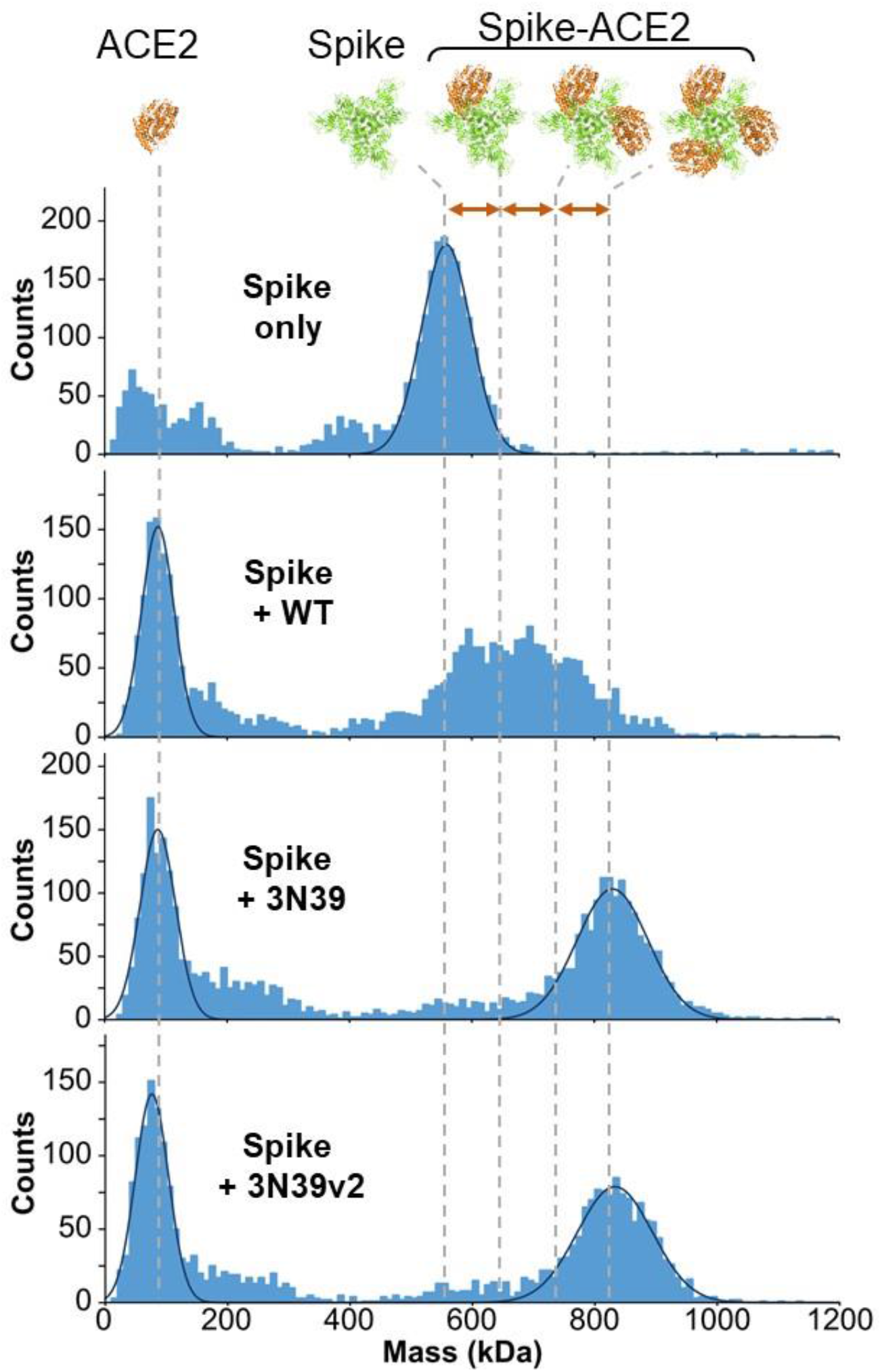
Mass photometry histograms of SARS-CoV-2 spike trimer in the absence or presence of ACE2 molecules. An average mass of spike trimer was measured to be ∼560 kDa (top panel). All ACE2 molecules added in excess to the mixture samples gave a peak with an average mass of ∼90 kDa. The spike protein complexed with WT ACE2 showed a broad mass distribution between 550∼850 kDa (the second panel). In contrast, the spike protein complexed with the 3N39 or 3N39v2 showed a monodisperse distribution with an average mass of ∼830 kDa (the third and bottom panels), which corresponds to the mass of the spike trimer bound by three ACE2 molecules.

**Fig S5.**
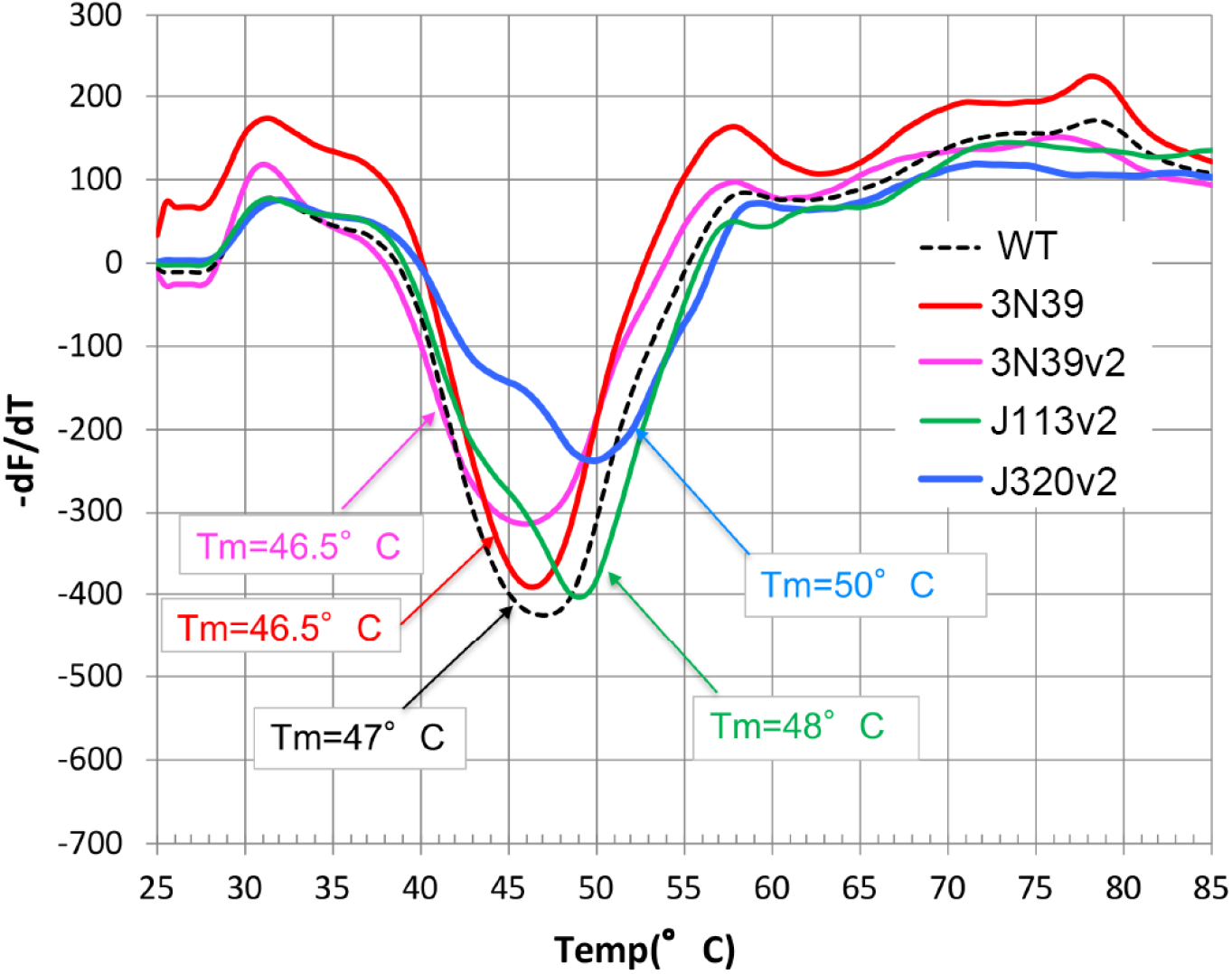
Thermostability of ACE2 mutants. Various versions of ACE2-His proteins were subjected to the differential scanning fluorimetry using SYPRO™ Orange as the probe dye. Denaturation curves were replotted for -*d*F/*d*T and the peak temperature was estimated to be the Tm for each mutant.

**Fig S6.**
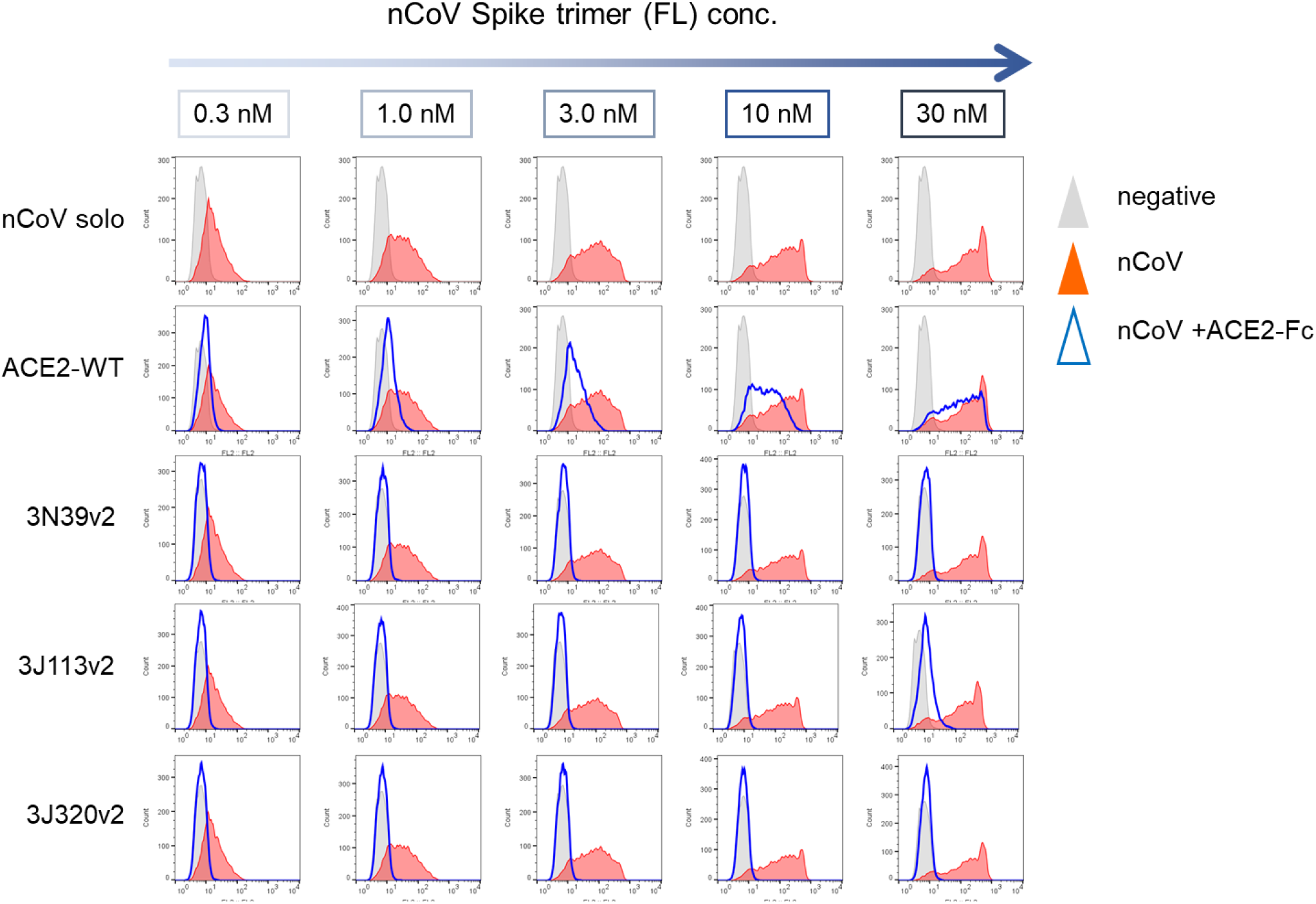
Neutralization of spike trimer by ACE2-Fcs. Neutralizing activity of ACE2-Fcs against soluble spike trimer binding to cell-surface ACE2 was evaluated in flow cytometry. Indicated concentration of spike trimer was incubated with 60 µg/ml (∼315 nM) ACE2-Fc proteins for 2h and then the mixture was reacted with ACE2-expressing Expi293F cells. Although WT ACE2-Fc can block binding of soluble spike protein to cells when the concentration of the spike was 0.3 nM, it could not outcompete spike at higher concentration. In contrast, complete (for 3N39v2 and J320v2) or near-complete (for J113v2) inhibition against 30 nM spike protein was achieved with the ACE2 mutants we have isolated, indicating the >100-fold increase in the blocking efficacy from the WT. The experiments were independently performed twice and similar results were obtained. One representative data were shown.

**Fig S7.**
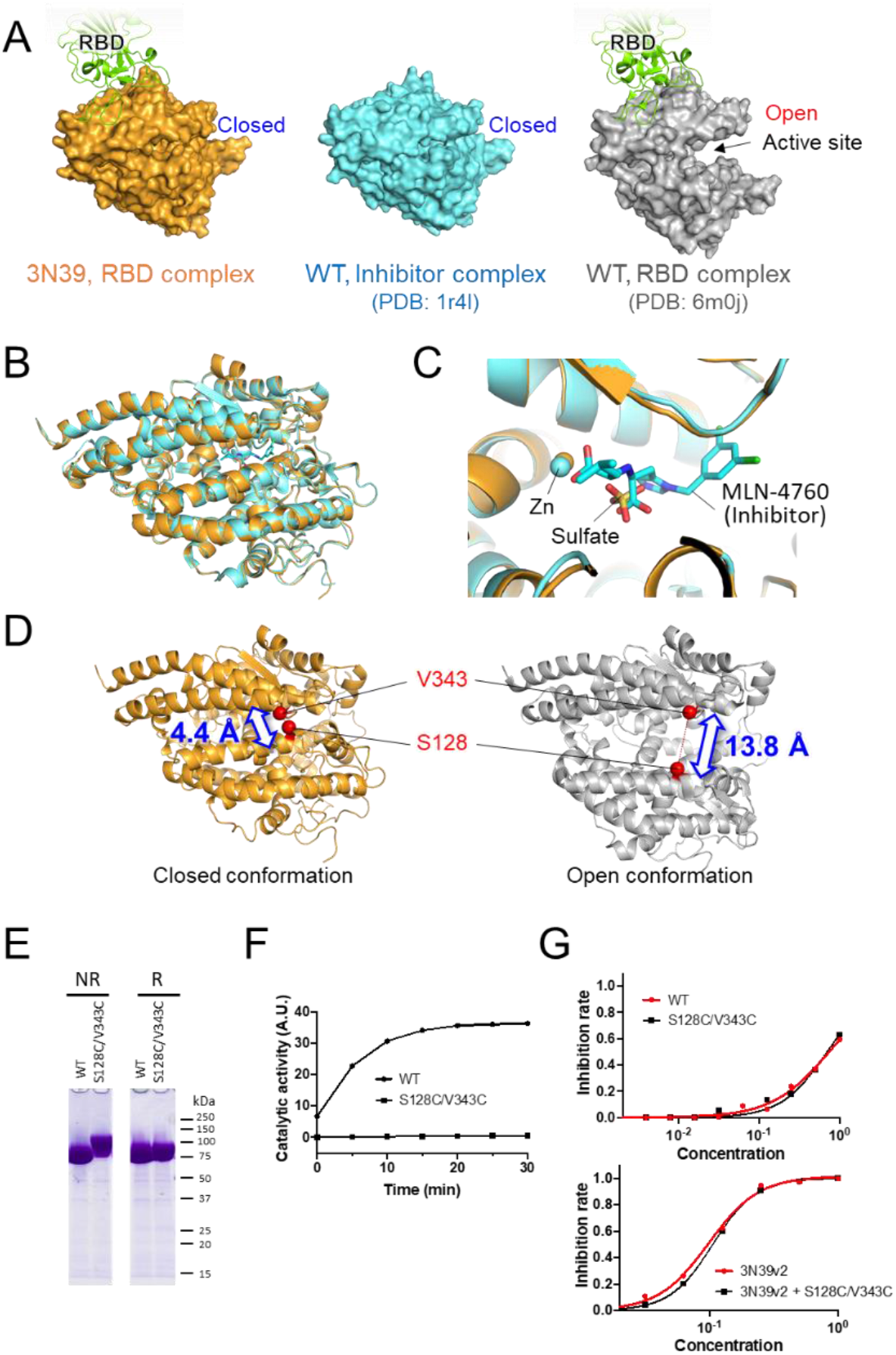
Open/closed conformations of ACE2. (**A**) Surface representations of the ACE2 structures. All ACE2 molecules are viewed from the same orientation (side view). The RBD-bound 3N39 (left) and the inhibitor-bound WT (center, 1r4l) adopt a closed conformation, while the RBD-bound WT (right, 6m0j) adopts an open conformation. (**B** and **C**) Superposition of the RBD-bound 3N39 (orange) and the inhibitor-bound WT (cyan, 1r4l) structures. Overall view and the expanded view of the enzymatic active site are provided in (B) and (C), respectively. The active site Zn ions (in both structures) are shown as sphere models, and a sulfate ion (in 3N39) and the inhibitor MLN-4760 (in 1r4l) are shown as stick models. (**D**) Comparison of the distances between C*β* atoms of S128 and V343 residues in the closed and open conformations. (**E**) SDS-PAGE analysis of WT and S128C/V343C mutant ACE2-His samples conducted under non-reducing (NR) and reducing (R) conditions. (**F**) Enzyme activity was assayed in the form of soluble ACE2-sfGFP by measuring Mca fluorescence resulting from hydrolysis of Mca-Ala-Pro-Lys(Dnp)-OH. (**G**) The RBD neutralization assay was performed in S128C/V343C mutant. The experiments were independently performed twice and similar results were obtained. One representative data were shown.

**Fig S8.**
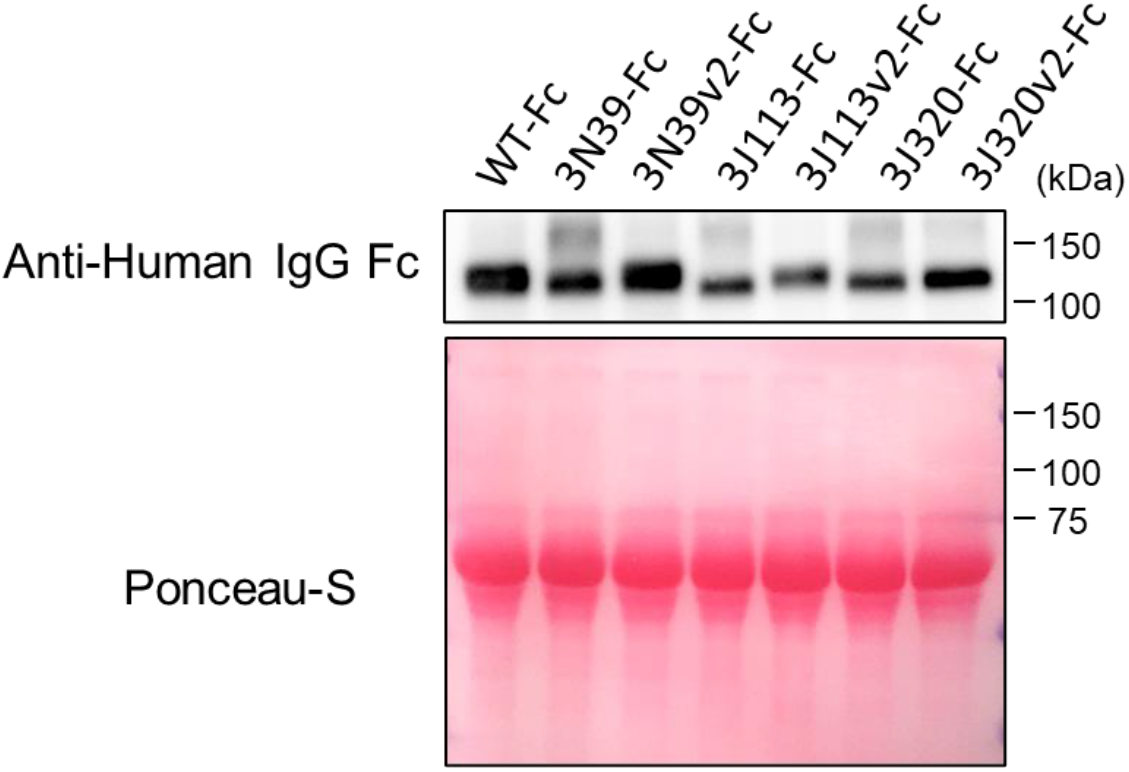
The yield of ACE2-Fc proteins. Western blot of cultured medium from each ACE2-Fc transfected 293T cells. The experiments were independently performed 3 times and similar results were obtained. One representative data were shown.

**Fig S9.**
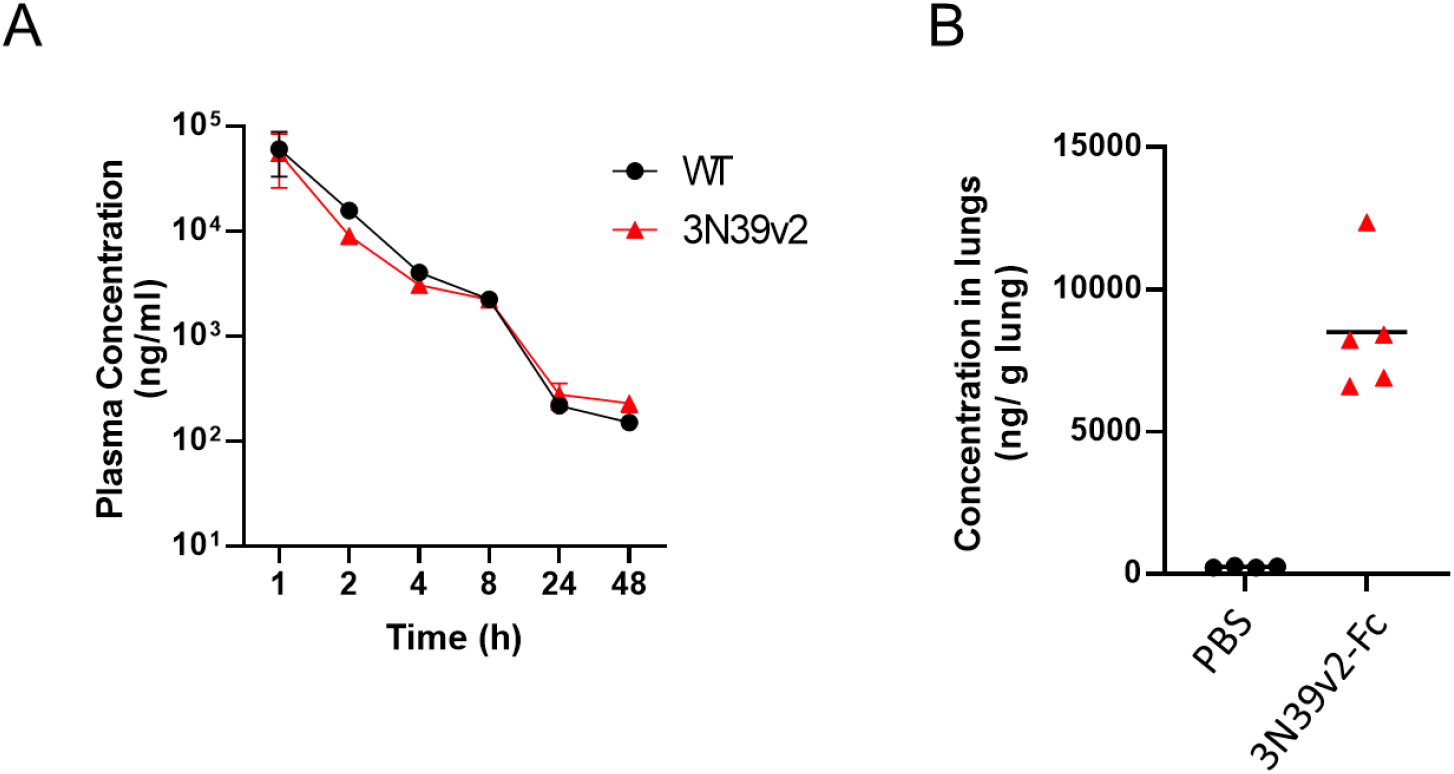
Pharmacokinetics of ACE2-Fc after single dose administration. Single dose of wild-type or 3N39v2-Fc (20mg/kg) was intraperitoneally administered in mice. Plasma concentration (**A**) in indicated time points and lung concentration (**B**) 2 hr after injection was analyzed by ELISA method. (A) Data are mean ± SD of n = 3 (WT) and 4 (3N39v2).

**Fig S10.**
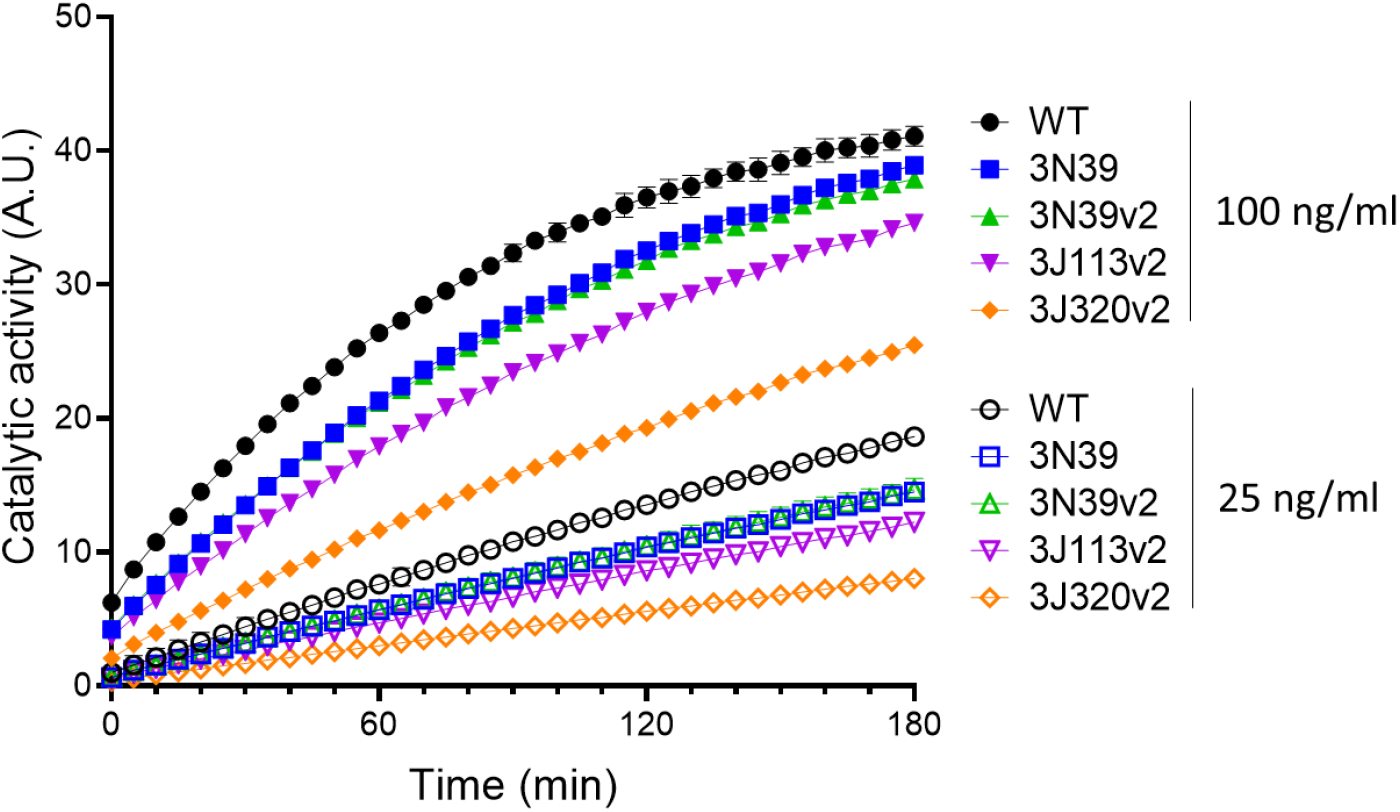
ACE2 catalytic activity of each mutants. Enzyme activity was assayed by measuring Mca fluorescence resulting from hydrolysis of Mca-Ala-Pro-Lys(Dnp)-OH. Fluorescence was recorded every 5 min. Data are mean ± SD of n = 3 technical replicates.

**Table S1.**
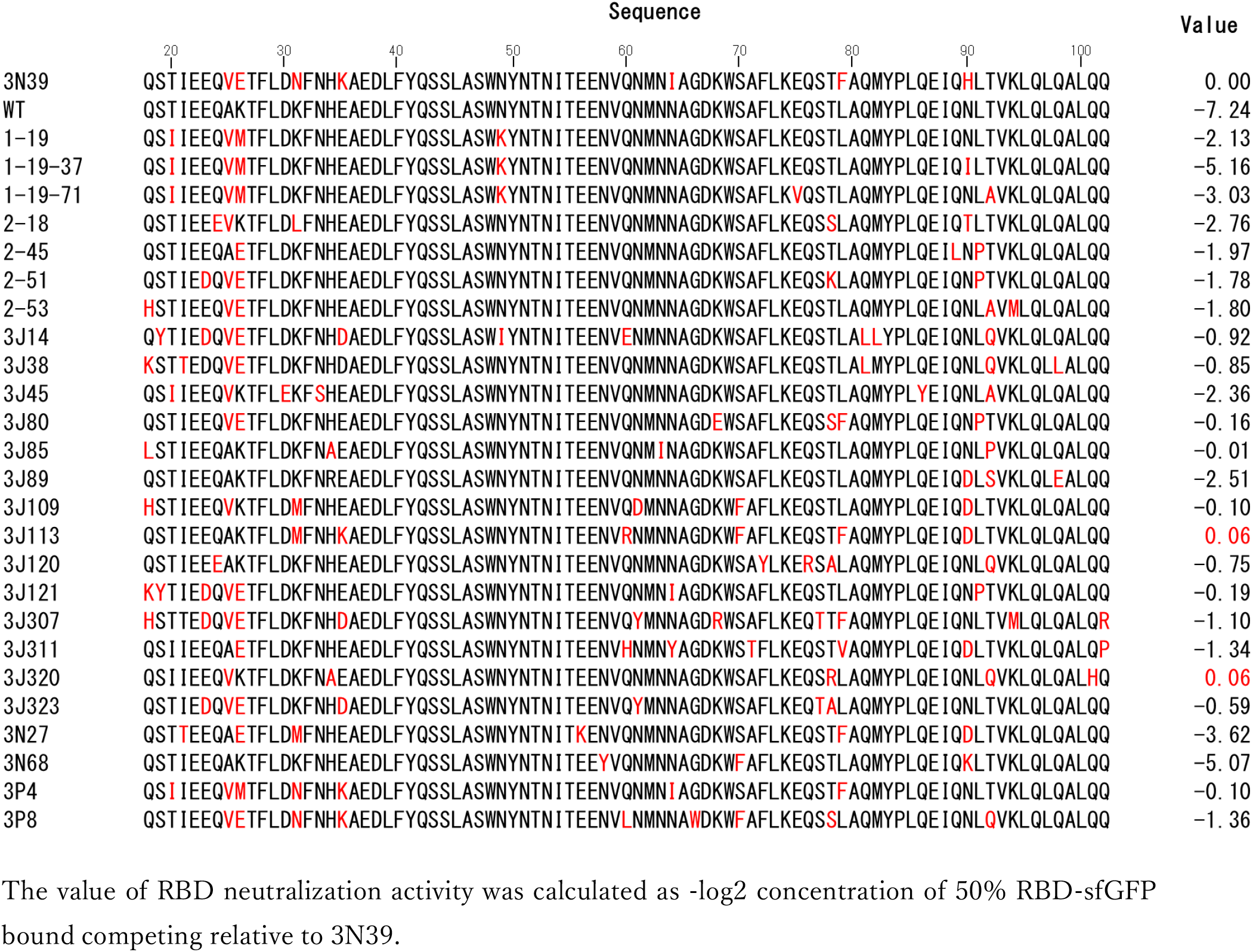
Amino acid sequence and RBD neutralization activity value of validated mutants.

**Table S2.** Data collection and refinement statistics

| 3N39 ACE2-RBD complex<br>(PDB: 7dmu) |  |
| --- | --- |
| **Data collection<sup>a</sup>** |  |
| Space group | *P4<sub>3</sub>* |
| Cell dimensions |  |
| *a, b, c* (Å) | 227.8, 227.8, 147.0 |
| Resolution (Å) | 48.13 - 3.20 (3.28 - 3.20)<sup>b</sup> |
| *R*<sub>merge</sub> | 0.29 (7.03) |
| *I* / $\sigma I$ | 9.38 (0.56) |
| CC1/2 (%) | 99.9 (29.3) |
| Completeness (%) | 99.9 (99.8) |
| Redundancy | 32.5 (28.9) |
| **Refinement** |  |
| Resolution (Å) | 47.91 – 3.20 |
| No. reflections | 123,205 |
| *R*<sub>work</sub> / *R*<sub>free</sub> (%) | 17.9/19.8 |
| No. atoms |  |
| Protein | 12,855 |
| Carbohydrates | 437 |
| Others | 14 |
| *B*-factors |  |
| Proteins | 132.9 |
| Carbohydrates | 209.0 |
| Others | 152.5 |
| R.m.s. deviations |  |
| Bond lengths (Å) | 0.007 |
| Bond angles (°) | 0.960 |

<sup>a</sup>Four datasets collected from a single crystal were merged.
<sup>b</sup>Values in parentheses are statistics of the highest-resolution shell.

MOVIE S1. Three-dimensional integrated CT images of the lungs from an uninfected hamster. The movie starts form front view.

MOVIE S2. Three-dimensional integrated CT images of the lungs from an infected hamster with the control-Fc treatment. Intact area is shown in white invertedly. The movie starts form front view.

MOVIE S3. Three-dimensional integrated CT images of the lungs from an infected hamster with the 3N39v2-Fc treatment. Intact area is shown in white invertedly. The movie starts form front view.

